# Revisiting tradeoffs between Rubisco kinetic parameters

**DOI:** 10.1101/470021

**Authors:** Avi I. Flamholz, Noam Prywes, Uri Moran, Dan Davidi, Yinon M. Bar-On, Luke M. Oltrogge, Rui Alves, David Savage, Ron Milo

## Abstract

Rubisco is the primary carboxylase of the Calvin cycle, the most abundant enzyme in the biosphere, and one of the best-characterized enzymes. Based on correlations between Rubisco kinetic parameters, it is widely posited that constraints embedded in the catalytic mechanism enforce tradeoffs between CO_2_-specificity, S_C/O_, and maximum carboxylation rate, *k*_cat,C_. However, the reasoning that established this view was based on data from ≈20 organisms. Here, we re-examine models of tradeoffs in Rubisco catalysis using a dataset from ≈300 organisms. Correlations between kinetic parameters are substantially attenuated in this larger dataset, with the inverse relationship between *k*_cat,C_ and S_C/O_ being a key example. Nonetheless, measured kinetic parameters display extremely limited variation, consistent with a view of Rubisco as a highly-constrained enzyme. More than 95% of *k*_cat,C_ values are between 1 and 10 s^−1^ and no measured k_cat,C_ exceeds 15 s^−1^. Similarly, S_C/O_ varies by only 30% among Form I Rubiscos and < 10% among C_3_ plant enzymes. Limited variation in S_C/O_ forces a strong positive correlation between the catalytic efficiencies (*k*_cat_/K_M_) for carboxylation and oxygenation, consistent with a model of Rubisco catalysis in which increasing the rate of CO_2_ addition to the enzyme-substrate complex requires an equal increase to the O_2_ addition rate. Altogether, these data suggest that Rubisco evolution is tightly constrained by the physicochemical limits of CO_2_/O_2_ discrimination.

## Introduction

Ribulose-1,5-Bisphosphate Carboxylase/Oxygenase (Rubisco) is the primary carboxylase of the Calvin-Benson-Bassham (CBB) cycle - the carbon fixation cycle responsible for growth throughout the green lineage and many other autotrophic taxa - and the ultimate source of nearly all carbon atoms entering the biosphere.^1^ Typically, 20-30% of total soluble protein in C_3_ plant leaves is Rubisco.^2^ As Rubisco is so highly expressed and plants are the dominant constituents of planetary biomass,^3^ it is often said that Rubisco is the most abundant enzyme on Earth. ^1,4^ Since Rubisco is ancient (> 2.5 billion years old), abundant, and remains central to biology, one might expect it to be exceptionally fast. But Rubisco is not fast.^5–8^ Typical central metabolic enzymes have a turnover number *k*_cat_ ≈ 80 s^−1^,^7^ but > 95% of Rubisco carboxylation *k*_cat,C_ values are between 1-10 s^−1^ and no measured *k*_cat,C_ values exceed 15 s^−1^.

In addition to relatively low *k*_cat,C_ values, Rubisco reacts with O_2_ in a process called oxygenation (Figure 1A). Although both carboxylation and oxygenation of the five-carbon substrate ribulose 1,5-bisphosphate (RuBP) are energetically favorable,^9^ carboxylation is the productive reaction for incorporating carbon from CO_2_ into precursors that generate biomass (Figure 1B). While it may play a role in sulfur, nitrogen, and energy metabolism,^10,11^ oxygenation is often considered counterproductive as it occupies Rubisco active sites and yields a product, 2-phosphoglycolate (2PG), that is not part of the CBB cycle and must be recycled through metabolically-expensive photorespiration at a partial loss of carbon.^12^ As such, oxygenation can substantially reduce the net rate of carboxylation by Rubisco, depending on CO_2_ and O_2_ concentrations and the kinetic parameters of the particular enzyme. There are at least four distinct Rubisco isoforms in nature,^13^ but all isoforms catalyze carboxylation and oxygenation of RuBP through the multistep mechanism described in Figures 1A and 1C.^14,15^ Even though many autotrophs depend on Rubisco carboxylation for growth, all known Rubiscos are relatively slow carboxylases and fail to exclude oxygenation.

**Figure 1:**
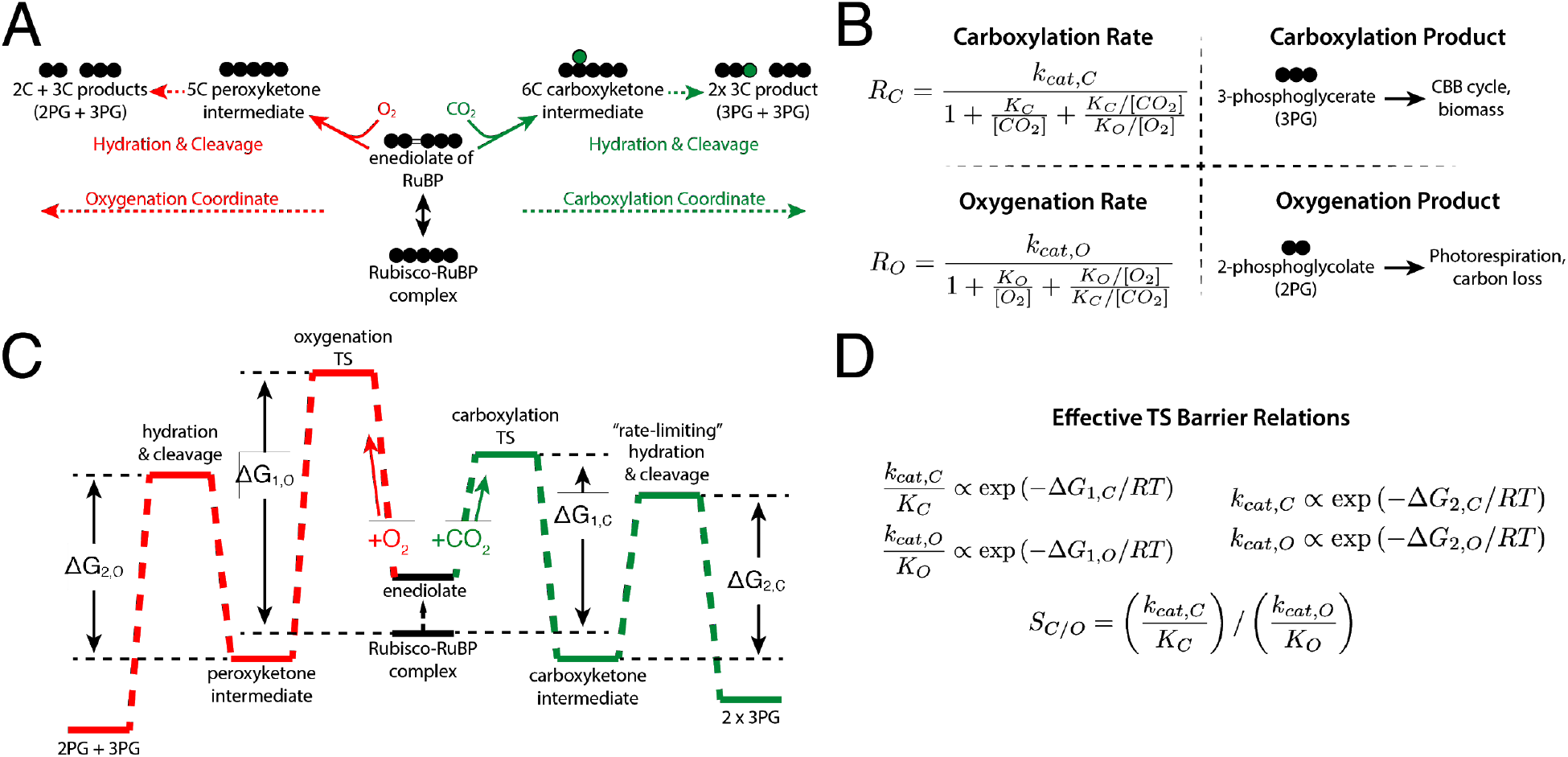
Description of the catalytic mechanism of Rubisco. The “middle-out” diagram in (A) shows the ordered mechanisms of carboxylation and oxygenation. Circles represent carbon atoms. RuBP is isomerized to an enediolate before carboxylation or oxygenation. Addition of CO_2_ or O_2_ to the enediolate of RuBP are considered irreversible as are the subsequent hydration and cleavage steps of carboxylation and oxygenation arms. (B) Carboxylation displays effective Michaelis-Menten kinetics (maximum catalytic rate k_cat,C_, half-maximum CO_2_ concentration K_M_ = K_C_) with competitive inhibition by O_2_ (assuming halfmaximum inhibitory O_2_ concentration K_i_ = K_O_). Carboxylation results in net addition of one carbon to the five-carbon RuBP, producing two 3PG molecules. 3PG is part of the CBB cycle and can therefore be used to continue the cycle and produce biomass. Oxygenation also displays effective Michaelis-Menten kinetics (k_cat,O_, K_M_ = K_O_, half-max inhibitory CO_2_ concentration K_I_ = K_C_). Oxygenation of RuBP produces one 3PG and one 2PG. Rates of carboxylation (R_C_) and oxygenation (R_O_) are calculated from kinetic parameters and the CO_2_ and O_2_ concentrations. The reaction coordinate diagram in (C) describes carboxylation and oxygenation as a function of two “effective” barriers.^6^ The first effective barrier includes enolization and gas addition while the second includes hydration and cleavage. (D) Given standard assumptions (SI), catalytic efficiencies (k_cat_/K_M_) are related to the height of the first effective barrier while k_cat_s are related to the second. The first barrier to oxygenation is drawn higher than for carboxylation because oxygenation is typically slower than carboxylation. Net reactions of RuBP carboxylation and oxygenation are both quite thermodynamically favorable.^9^

The fastest-carboxylating Rubisco observed (at 25 °C) is from the cyanobacterium *S. elongatus* PCC 7942.^16^ This enzyme has a maximum per-active site carboxylation rate (*k_cat,C_*) of 14 s^−1^. However, because present-day atmosphere contains abundant O_2_ and relatively little CO_2_ (≈21% O_2_, ≈0.04% CO_2_), PCC 7942 Rubisco carboxylates at a rate 20-fold below maximum in ambient conditions (R_C_ ≈ 0.7 s^−1^ per active site, rate law in Figure 1A). Due to its relatively low CO_2_-specificity, PCC 7942 Rubisco will also oxygenate RuBP appreciably in ambient conditions (R_O_ ≈ 0.3 s^−1^), necessitating substantial photorespiratory flux to recycle 2PG. As downstream processing of 2PG by the C_2_ photorespiratory pathway leads to the loss of one carbon for every two 2PG,^11,12^ every two oxygenations “undoes” a carboxylation. In ambient air, therefore, the net rate of carboxylation by PCC 7942 Rubisco would be *f* = R_C_ − R_O_/2 ≈ 0.6 s^−1^, or ≈4% of *k_cat,C_*. Given the kinetics of PCC 7942 Rubisco, it is not surprising that all known cyanobacteria use a CO_2_-concentrating mechanism to ensure Rubisco functions in a CO_2_-rich environment. Elevated CO_2_ ensures that oxygenation is competitively inhibited and that carboxylation proceeds at near-maximum rate.^17^ Thirty fold enrichment of CO_2_ above ambient increases the carboxylation rate of PCC 7942 Rubisco to 8.9 s^−1^ and suppresses oxygenation to ≈0.14 s^−1^, giving a net carboxylation rate *f* = 8.8 s^−1^ per active site (≈60% of *k_cat,C_*).

For comparison, the Rubisco from spinach leaves (*S. oleracea*) is characteristic of plant Rubiscos in having lower k_cat,C_ ≈ 3 s^−1^ and much greater CO_2_-affinity than the *S. elongatus* enzyme (spinach half-maximum CO_2_ concentration K_C_ ≈ 12 μM, PCC 7942 K_C_ ≈ 170 μM). As a result, the spinach enzyme outperforms the cyanobacterial one in ambient air, with R_C_ ≈ 1.2 s^−1^, R_O_ ≈ 0.4 s^−1^, and *f* ≈ 1 s^−1^. Spinach is a C_3_ plant, meaning it does not have a CO_2_-concentrating mechanism, which may explain why it employs a slow-but-specific Rubisco. Still, most central metabolic enzymes catalyze far more than 1 reaction per second,^7^ leading many to wonder if Rubisco catalysis could be improved. Improved Rubisco carboxylation might increase C_3_ crop yields, ^18,19^ but a substantially improved enzyme has evaded bioengineers for decades.^20^ The repeated evolution of diverse CO_2_ concentrating mechanisms, which modulate the catalytic environment rather than Rubisco itself, raises further doubts about whether Rubisco catalysis can be strictly improved.^21^

Various nomenclature has been used to describe the kinetics of Rubisco carboxylation and oxygenation since its discovery in the 1950s.^5,6,22,23^ Here we use k_cat,C_ and k_cat,O_ to denote turnover numbers (maximum rates per active site, s^−1^ units) for carboxylation and oxygenation respectively. K_C_ and K_O_ denote the Michaelis constants (half-saturation concentrations in μM units) for carboxylation and oxygenation. The specificity factor S_C/O_ = (k_cat,C_/K_C_) / (k_cat,O_/K_O_) is a unitless measure of the relative preference for CO_2_ over O_2_ (Figure 1D). Since S_C/O_ relates only to the ratio of kinetic parameters, higher S_C/O_ does not necessarily imply higher carboxylation rates. Rather, absolute carboxylation and oxygenation rates depend on CO_2_ and O_2_ concentrations (Figure 1B) which can vary between organisms and environments (SI).

As data on bacterial, archaeal and plant carboxylases has accumulated over the decades, many researchers have noted that fast-carboxylating Rubiscos are typically less CO_2_-specific.^24–26^ In other words, Rubiscos with high k_cat,C_ were observed to have lower S_C/O_ due either to lower CO_2_-affinity (high K_C_) or more efficient oxygenation (higher k_cat,O_/K_O_). Negative correlation between k_cat,C_ and S_C/O_ is often cited to motivate the idea that a tradeoff between carboxylation rate and specificity constrains Rubisco evolution.^5,6,26,27^

It is worth pausing to clarify the concepts of “tradeoff,” “constraint,” and “correlation” (Figure 2). Correlation indicates an apparent linear (or log-log, etc.) relationship between two kinetic parameters. Correlations between enzyme kinetic parameters can result from a “tradeoff” due to two distinct kinds of underlying constraints (Figure 2).^6^ In the “mechanistic coupling” scenario, the enzymatic mechanism forces a strict quantitative relationship between two kinetic parameters such that varying one forces the other to vary in a defined manner. This results in a situation where the value of one parameter strictly determines the other and vice versa (i.e. an “equality constraint,” Figure 2A). This could arise for Rubisco, for example, if a single catalytic step (e.g. enolization of RuBP, Figure 1A) determines the rates of CO_2_ and O_2_ entry both.

**Figure 2:**
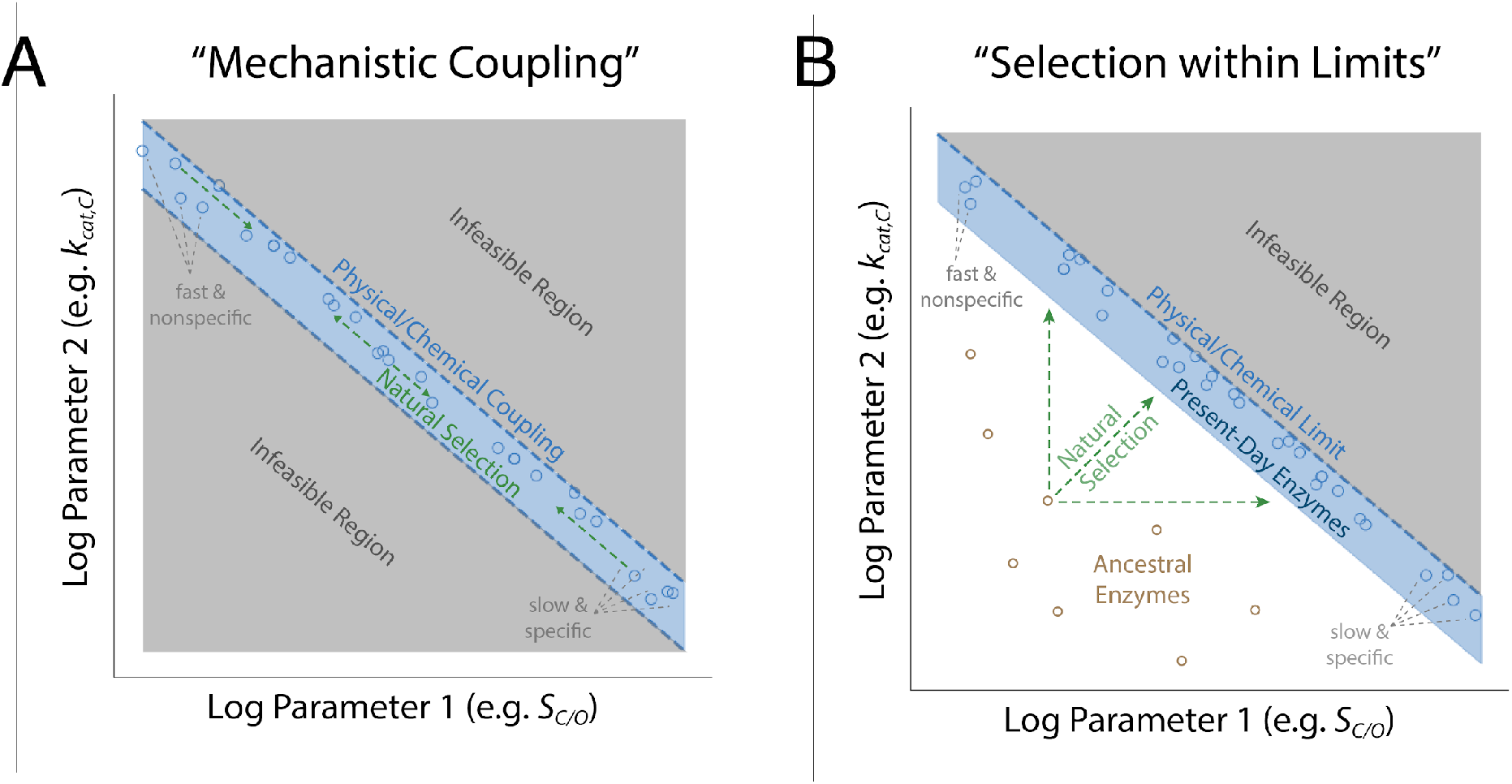
Scenarios that produce strong correlations between enzyme kinetic parameters. As the log of the kinetic parameters are linearly related to energy barriers, linear energetic tradeoffs should manifest as log-log correlations between kinetic parameters (power laws). Panel (A) describes a situation in which two kinetic parameters are inextricably linked by the enzyme mechanism, diagrammed here as negative coupling between k_cat,C_ and S_C/O_ as an example. These couplings take the form of “equality constraints” where one parameter determines the other within measurement error. Correlation is expected so long as diverse enzymes are measured. In (A) selection moves enzymes along the blue curve, but cannot produce enzymes off the curve (grey) because they are infeasible. Panel (B) diagrams an alternative scenario in which the enzyme mechanism imposes an upper limit on two parameters (an inequality constraint). In the “selection within limits” scenario, effective selection is required for correlation to emerge because sub-optimal enzymes (e.g. ancestral sequences) are feasible. In the examples plotted, different environmental CO_2_ and O_2_ concentrations should select for different combinations of rate (k_cat,C_) and affinity (S_C/O_), resulting in present-day enzymes occupying distinct regions of the plots in (A) and (B).

In the “selection within limits” model (Figure 2B), in contrast, the catalytic mechanism imposes an upper limit on kinetic parameters, i.e. an inequality constraint. A clear correlation between parameters will emerge only if there is sufficient selection to reach the boundary. To highlight the difference between these models, consider the kinetics of ancestral enzymes. In the “mechanistic coupling” model, kinetic parameters of ancestors should lie along the same curve as present-day enzymes because the grey regions off the curve are disallowed. Selection could act by moving enzymes along the line of mechanistic coupling - e.g. from a region of high selectivity and low rate towards higher rate and lower selectivity (Figure 2A). According to the “selection within limits” model, in contrast, ancestral enzymes can lie beneath the upper limit determined by the catalytic mechanism (Figure 2B). This second model requires selection to produce a situation where the kinetics of present-day Rubiscos extracted from various organisms trace out a curve determined by the upper limit enforced by the mechanism.^28^

Previous research advanced two distinct families of mechanistic models to explain correlations between Rubisco kinetic parameters.^5,6^ The first model, which we term “k_cat,C_-K_C_ coupling,” hypothesizes a tradeoff between the rate and affinity of carboxylation that leads to a negative correlation between k_cat,C_ and S_C/O_ (Figure S2).^5^ A second model, which was advanced in a study which the last author of this work participated in,^6^ hypothesizes that multiple tradeoffs constrain Rubisco such that kinetic parameters can only vary along a one dimensional curve. In addition to k_cat,C_-K_C_ coupling, this work hypothesized a tradeoff between catalytic efficiencies for carboxylation and oxygenation (coupling k_cat,C_/K_C_ and k_cat,O_/K_O_) wherein improving carboxylation efficiency also improves oxygenation efficiency.

These mechanistic models are substantively different. Though both models imply limitations on the concurrent improvement of k_cat,C_ and S_C/O_, “k_cat,C_-K_C_ coupling” relates only to carboxylation kinetics, leaving the possibility that oxygenation kinetics are unconstrained. Coupling between k_cat,C_/K_C_ and k_cat,O_/K_O_, in contrast, relates to both reaction pathways. While these models appeal to physical and chemical intuition, they are based on data from only ≈20 organisms. Moreover, “mechanistic coupling” and “selection within limits” could plausibly underly either model (Figure 2).^6^

Here we take advantage of the accumulation of new data to revisit correlations and tradeoffs between Rubisco kinetic parameters. We collected and curated literature measurements of ≈300 Rubiscos. Though diverse organisms are represented, the Form I Rubiscos of C_3_ plants make up the bulk of the data (>80%, Figure 3A). Most previously-reported correlations between Rubisco kinetic parameters are substantially attenuated in this dataset, with the negative correlation between k_cat,C_ and specificity S_C/O_ being a key example. Weakened k_cat,C_-S_C/O_ and k_cat,C_-K_C_ correlations imply that these parameters are not straightforwardly mechanistically coupled, suggesting that models of k_cat,C_-K_C_ coupling should be revisited in future experiments. Overall, weakened correlations call into question previous claims that (i) Rubisco kinetics are constrained to evolve on a one-dimensional line and (ii) natural Rubiscos are optimized to suit environmental CO_2_ and O_2_ concentrations.^5,6^

**Figure 3:**
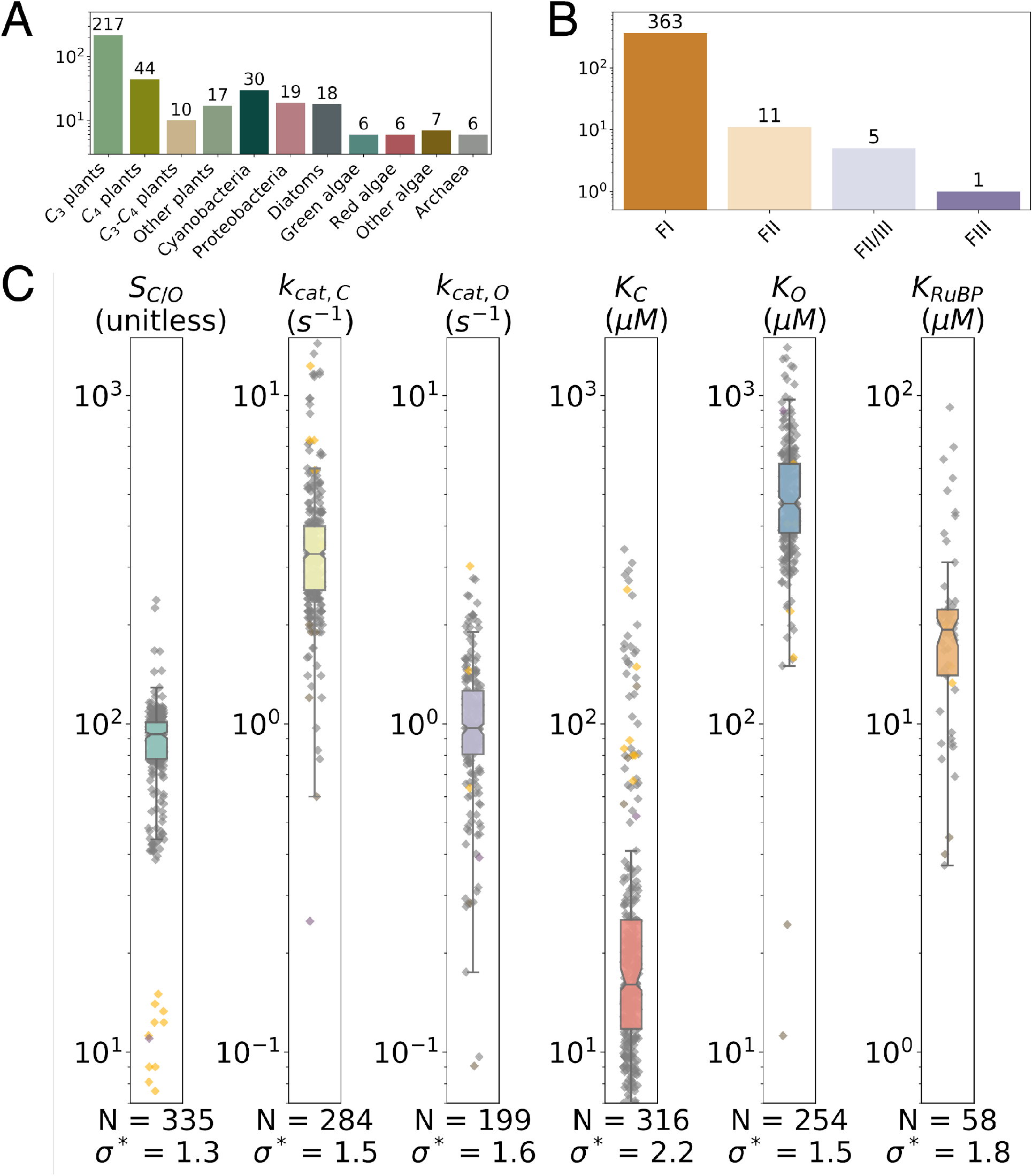
Summary of the full extended dataset. We collected measurements of Rubisco kinetic parameters from a variety of organisms (A) representing four classes of Rubisco isoforms (B). Form I enzymes from plants, cyanobacteria and algae make up the bulk of the data (A-B). (C) Rubisco kinetic parameters display narrow dynamic range. The box-plot and grey points describe the distribution of Form I Rubiscos while Form II Rubiscos are in yellow. Colored boxes give the range of the central 50% of FI values and the notch indicates the median. N is the number values and σ* gives the geometric standard deviation of Form I data. σ* < 3 for all parameters, meaning a single standard deviation varies over less than threefold. All data are from wild-type Rubiscos measured at 25 °C near pH 8. More detailed histograms are given in Figure S4.

Despite weakened correlations, Rubisco kinetic parameters display extremely limited variation. k_cat,C_ varies by only 50% among Form I Rubiscos, and S_C/O_ varies even less than that (≈30%, Figure 3C). Limited variation in S_C/O_ forces a strong positive power-law correlation between the catalytic efficiencies for carboxylation (k_cat,C_/K_C_) and oxygenation (k_cat,O_/K_O_).^6^ We propose a simple model of mechanistic coupling that explains how constraints on the Rubisco mechanism could restrict variation in S_C/O_. In this model, variation in catalytic efficiency (k_cat,C_/K_C_ and k_cat,O_/K_O_) derives solely from gating substrate access to the active site complex, which could help explain why Rubisco has been so recalcitrant to improvement by mutagenesis and rational engineering.

## Materials and Methods

### Data collection and curation

We reviewed the literature to find Rubisco kinetic data measured at 25 °C and near pH 8. Ultimately 61 primary literature studies were included, yielding 335 S_C/O_, 284 k_cat,C_, 316 K_C_, and 254 K_o_ values for Rubiscos from 304 distinct organisms (Datasets S1 and S2). We also recorded 58 measurements of the Michaelis constant for RuBP (K_RuBP_). Experimental error was recorded for all of these values (when reported) along with the pH, temperature and other metadata. Data was filtered as described in SI. k_cat,O_ is usually not measured directly,^29^ but is rather inferred as k_cat,O_ = (k_cat,C_/K_C_) / (S_C/O_/K_O_). We assumed that experimental error is normally distributed and used 10^4^-fold bootstrapping to estimate 199 k_cat,O_ values and 95% confidence intervals thereof. We used an identical procedure to estimate k_cat,C_/K_C_ and k_cat,O_/K_O_ and confidence intervals thereof (SI). Altogether, we were able to calculate 274 k_cat,C_/K_C_ and 199 k_cat,O_/K_O_ values. Datasets S1 and S2 provide all source and inferred data respectively.

### Fitting power laws

Certain model Rubiscos are measured frequently. For example, we found 12 independent measurements of the spinach Rubisco. In such cases, the median measured value was used to avoid bias in correlation and regression analyses. In contrast to textbook examples with one independent and one dependent variable, there is experimental error associated with both variables in all scatter plots shown here (e.g. plotting *k_cat,C_* against *K_C_* in Figure 5B). As such we used total least squares linear regression in log scale to fit relationships between Rubisco parameters. Because R^2^ values of total least squares fits do not convey the explained fraction of Y axis variance, they are challenging to interpret. We instead report the degree of correlation as Pearson R values of log-transformed values. Bootstrapping was used to determine 95% confidence intervals for the Pearson correlation coefficient, power-law exponents and prefactors (i.e. the slopes and intercepts of linear fits in log-log scale). In each iteration of the bootstrap, data were subsampled to 90% with replacement. Total least squares regression was applied to each subsample and the procedure was repeated 10^4^ times to determine 95% confidence intervals. Python source code is available at github.com/flamholz/rubisco.

## Results

### An extended dataset of Rubisco kinetic parameters

To augment existing data, we collected literature data on ≈300 Rubiscos including representatives of clades and physiologies that had been poorly represented in earlier datasets e.g. diatoms, ferns, CAM plants and anaerobic bacteria (Figure 3A). We collected kinetic parameters associated with carboxylation and oxygenation - S, K_C_, k_cat,C_, K_O_ and k_cat,O_ - as well as measurements of the RuBP Michaelis constant (half-maximum RuBP concentration, K_RuBP_) and experimental uncertainty for all values where available. All data considered below were measured at 25 °C and near pH 8 to ensure that measured values are comparable (SI). Notably, Rubisco assays are challenging to perform and variation in measurements across labs is expected. Some of the spread in the data may come from systematic differences between labs and assay methods. Rubisco activation state, for example, may differ between methods and preparations.^15^ Though we cannot resolve this issue here, we were careful to review each study’s methods, document a small number of problematic measurements, and record experimental error when reported (Dataset S1).

The resulting dataset contains Rubisco measurements from a total of 304 distinct species, including 335 S_C/O_ values, 284 k_cat,C_ values, 316 K_C_ values, 254 K_O_ values, and 199 k_cat,O_ values (Figure 3B). k_cat,O_ values are rarely measured directly (SI) and are typically inferred as k_cat,O_ = (k_cat,C_/K_C_) / (S_C/O_/K_O_).^29^ The Michaelis constant for RuBP (K_RuBP_) is measured infrequently and only 58 values were extracted. We were able to estimate catalytic efficiencies for carboxylation (k_cat,C_/K_C_) in 274 cases and for oxygenation (k_cat,O_/K_O_) in 199 cases (Methods). Though the data include measurements of some Form II, III and II/III Rubiscos, they remain highly focused on the Form I Rubiscos found in cyanobacteria, diatoms, algae and higher plants, which make up > 95% of the dataset (Figure 3B). As such, we focus here on the kinetic parameters of Form I Rubiscos, (abbreviated FI Rubisco).

Rubisco kinetic parameters display very narrow dynamic range (Figure 3C). The geometric standard deviation (*σ*\*) expresses multiplicative variability in the dataset and is well-below one order-of-magnitude (*σ*\* ≪ 10) for all parameters. Rubisco displays particularly low variation in *k_cat,C_* (*σ*\* = 1.5) as compared to other enzymes for which 20 or more *k_cat_* measurements are available (median *σ*\* ≈ 7, Figure S5). Specificity S_C/O_ displays the least variation of all parameters (*σ*\* = 1.3). This is due in part to overrepresentation of C_3_ plants in the dataset, which occupy a narrow range of S_C/O_ ≈ 80-120. Nonetheless, measurements of S_C/O_ for FI and FII enzymes are clearly distinct, with values ranging from and 7-15 for FII measurements and ≈50-200 for FI (Figure 3C).

### Energetic tradeoffs tend to produce power law correlations

All kinetic parameters (S_C/O_, k_cat,C_, K_C_, k_cat,O_ and K_O_) are mathematically related to the microscopic rate constants of the Rubisco mechanism (SI). Given common assumptions about irreversible and rate limiting steps (SI), this multi-step mechanism can be simplified so that logarithms of measured kinetic parameters are proportional to effective transition state barriers (Figure 1C-D, SI). As such, correlations between kinetic parameters will emerge if effective transition state barriers vary together (Figure 2). If, for example, lowering the effective transition state barrier to CO_2_ addition (ΔG_1,C_) requires an increase to the effective barrier to the subsequent hydration and cleavage steps of carboxylation (Δ_G2,C_), then we should observe a negative linear correlation *ΔG*_1,*C*_ ∝ −*ΔG*_2,*C*_. Since k_cat,C_/K_C_ is related to the first effective carboxylation barrier (*k_cat,C_*/*K_C_* ∝*exp* (−*ΔG*_1,*C*_/*RT*)) and k_cat,C_ to the second (*k_cat,C_* ∝*exp* (−*ΔG*_2,*C*_/*RT*)), linear correlation between transition state barrier heights translates to log-scale correlation between kinetic parameters such that *ln* (*k_cat,C_*/*K_C_*) ∝ − *ln* (*k_cat,C_*). These relationships are known as power laws and motivate us and others to investigate the kinetic parameters on a log-log scale.

We expect to observe strong power-law correlations between pairs of kinetic parameters in two cases. Either (i) the associated energy barriers co-vary because they are linked by the enzymatic mechanism (“mechanistic coupling,” Figure 2A) or (ii) the mechanism imposes an upper bound on the sum (or difference) of two barrier heights. In case (ii), strong selection favors the emergence of enzymes at-or-near the imposed limit (“selection within limits,” Figure 2B). As Rubisco is the central enzyme of photoautotrophic growth, it likely evolved under selection pressure towards maximizing the net rate of carboxylation in each host, and so either of these scenarios is plausible *a priori*. Notably, host physiology and growth environments can affect the catalytic environment: Rubiscos from different organisms will experience different temperature, pH and prevailing CO_2_ and O_2_ concentrations due, for example, to an anaerobic host or a CO_2_ concentrating mechanism enriching CO_2_.^6^ Different conditions should favor different combinations of kinetic parameters (Figure 2).

### Correlations between kinetic parameters of Form I Rubiscos

We performed a correlation analysis to investigate relationships between kinetic parameters of Form I Rubiscos. Figure 4 gives log-scale Pearson correlations between parameters that are measured directly: k_cat,C_, K_C_, K_O_, S_C/O_ and K_RuBP_. Linear scale correlations are reported in Figure S7.

**Figure 4:**
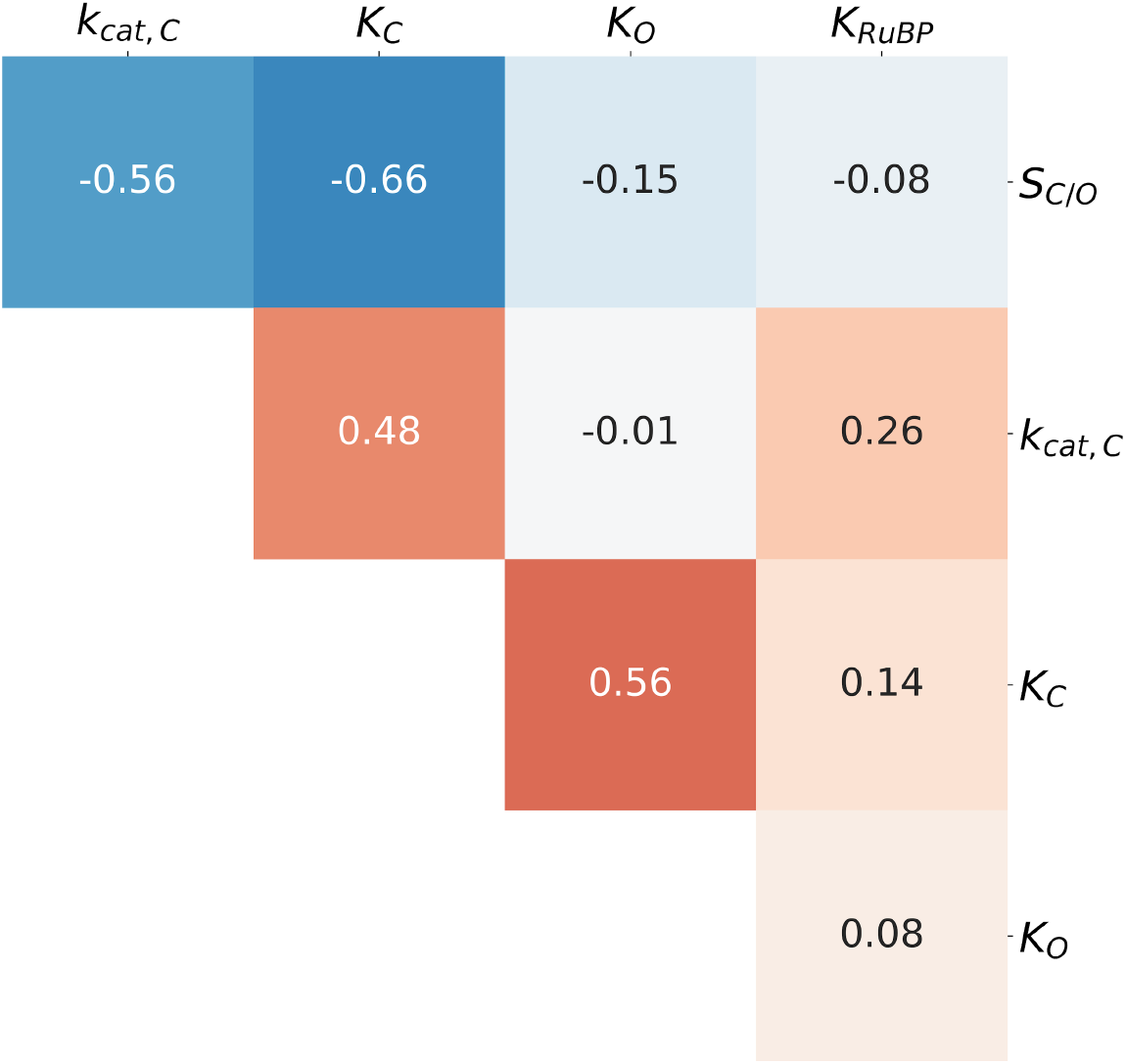
Correlations between measured kinetic parameters are attenuated by the addition of new data. The figure gives Pearson correlations (R) between pairs of log-transformed kinetic parameters of Form I Rubiscos. When multiple measurements of the same enzyme were available, the median value was used (Methods). S_C/O_-K_C_, S_C/O_-k_cat,C_, and K_C_-k_cat,C_ correlations are of particular interest because they were highlighted in previous works, which found R = 0.8-0.95. None of these pairs have R values exceeding 0.7 in the extended dataset.

Overall, correlations are weaker in the extended dataset than documented in previous studies of smaller datasets.^5,6^ Nonetheless, we observed modestly strong, statistically significant correlations between k_cat,C_ and S_C/O_ (R = −0.56, *p* < 10^−10^), k_cat,C_ and K_C_ (R = 0.48, *p* < 10^−10^), K_C_ and S_C/O_ (R = −0.66, *p* < 10^−10^), and K_C_ and K_O_ (R = 0.56, *p* < 10^−10^). Since Rubisco kinetic parameters are mathematically interrelated through the microscopic mechanism as it is commonly understood, some level of correlation is expected. For example, when we derive expressions for k_cat,C_ and K_C_ from the Rubisco mechanism, they share common factors that should produce some correlation even in the absence of underlying coupling (SI). Similarly, S_C/O_ is defined as (k_cat,C_/K_C_) / (k_cat,O_/K_O_) and could correlate negatively with K_C_ for this reason. Because modest correlation is expected irrespective of underlying tradeoffs, the correlations in Figure 4 do not necessarily support any particular tradeoff model.

Correlations between k_cat,C_ and S_C/O_ as well as k_cat,C_ and K_C_ were previously highlighted to support particular mechanistic tradeoff models.^5,6^ However, these correlations are substantially attenuated by the addition of new data (Figures 4 and 5). Plotting k_cat,C_ against S_C/O_ (Figure 5A) shows that these parameters are modestly correlated, with R ≈ 0.6 as compared to R ≈ 0.9 in previous analyses.^5,6^ Figure 5A also highlights the extremely limited and stereotyped variation in S_C/O_ where Rubiscos from organisms sharing a particular physiology (e.g. C_3_ plants or cyanobacteria) occupy a very narrow range of S_C/O_ values. Multiplicative standard deviations (*σ*\*) are less than 1.25 in all cases. Plotting k_cat,C_ against K_C_ (Figure 5B) shows that this correlation is also weakened, with R ≈ 0.5 as compared to ≈ 0.9 previously.^6^ We interpret weakened correlations as evidence that previously-proposed tradeoff models should be revisited. We therefore proceed to evaluate the correlations predicted by specific tradeoff models, with an eye towards understanding the restricted variation in S_C/O_ shown in Figure 5A.

**Figure 5:**
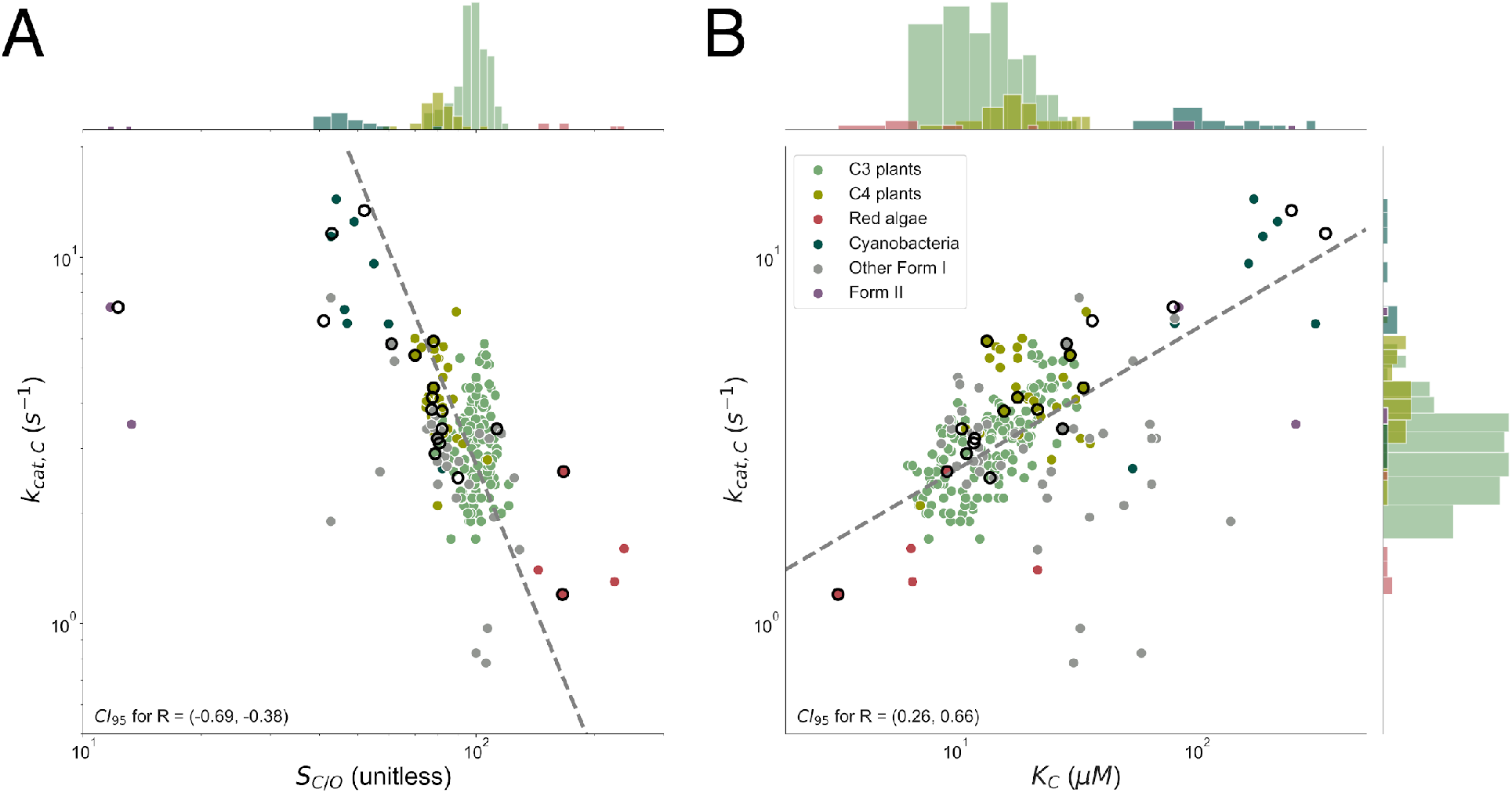
Focal correlations of previous analyses are not robust to new data. Points with black outlines are from Savir et al. 2010 and dashed grey lines represent the best fit to FI Rubisco data. Histograms for k_cat,C_, S_C/O_ and K_C_ are plotted on parallel axes. Panel (A) plots k_cat,C_ against S_C/O_. k_cat,C_ and S_C/O_ correlate with R ≈ −0.6 among FI Rubiscos as compared to ≈ 0.9 previously. ^5,6^ 95% confidence intervals are (−4.0, −2.0) for the fit exponent and (3×10^4^, 2×10^8^) for the prefactor (slope and intercept in log-log scale respectively) indicating that the form of k_cat,C_-S_C/O_ correlation is very uncertain. Notably, S_C/O_ displays very limited variation overall and especially within physiological groupings with sufficient data. Median S_C/O_ values are 177 for red algae (σ* = 1.2, N=6), 98 for C_3_ plants (σ* = 1.1, N=162), 80 for C_4_ plants (σ* = 1.1, N=35), and 48 for cyanobacteria (σ* = 1.1, N=16). Panel (B) plots k_cat,C_ against K_C_. Here, R ≈ 0.5 as compared to ≈ 0.9 previously.^6^ This fit is more robust, with 95% confidence intervals of (0.3, 0.5) and (0.8, 1.5) for the fit exponent and prefactor respectively.

### Re-evaluation of Proposed Tradeoff Models

Two distinct mechanistic tradeoff models have been advanced.^5,6^ The first model, which we term k_cat,C_-K_C_ coupling, posits that increased specificity towards CO_2_ necessitates a slower maximum carboxylation rate, k_cat,C_.^5,6^ It was proposed that this tradeoff is due to stabilization of the first carboxylation transition state (TS).^5^ Under this model, a stable Rubisco-TS complex produces high CO_2_-specificity but slows the subsequent carboxylation steps and limits k_cat,C_ (Figure S2). This proposal can be cast in energetic terms by relating the measured catalytic parameters to effective transition state barrier heights (Figure 1D, SI). This model can be construed in energetic terms as follows: lowering the effective barrier to CO_2_ addition (ΔG_1,C_ in Figure 6A) will make Rubisco more CO_2_-specific even if oxygenation kinetics remain unchanged.^6^ k_cat,C_-K_C_ coupling posits a negative coupling between CO_2_ addition and the subsequent carboxylation steps of hydration and bond cleavage (effective barrier height ΔG_2,C_ diagrammed in Figure 6A). Therefore, the energetic interpretation of this model predicts a negative correlation between ΔG_1,C_ and ΔG_2,C_ and, as a result, a negative power-law correlation between k_cat,C_ and k_cat,C_/K_C_.^6^

**Figure 6:**
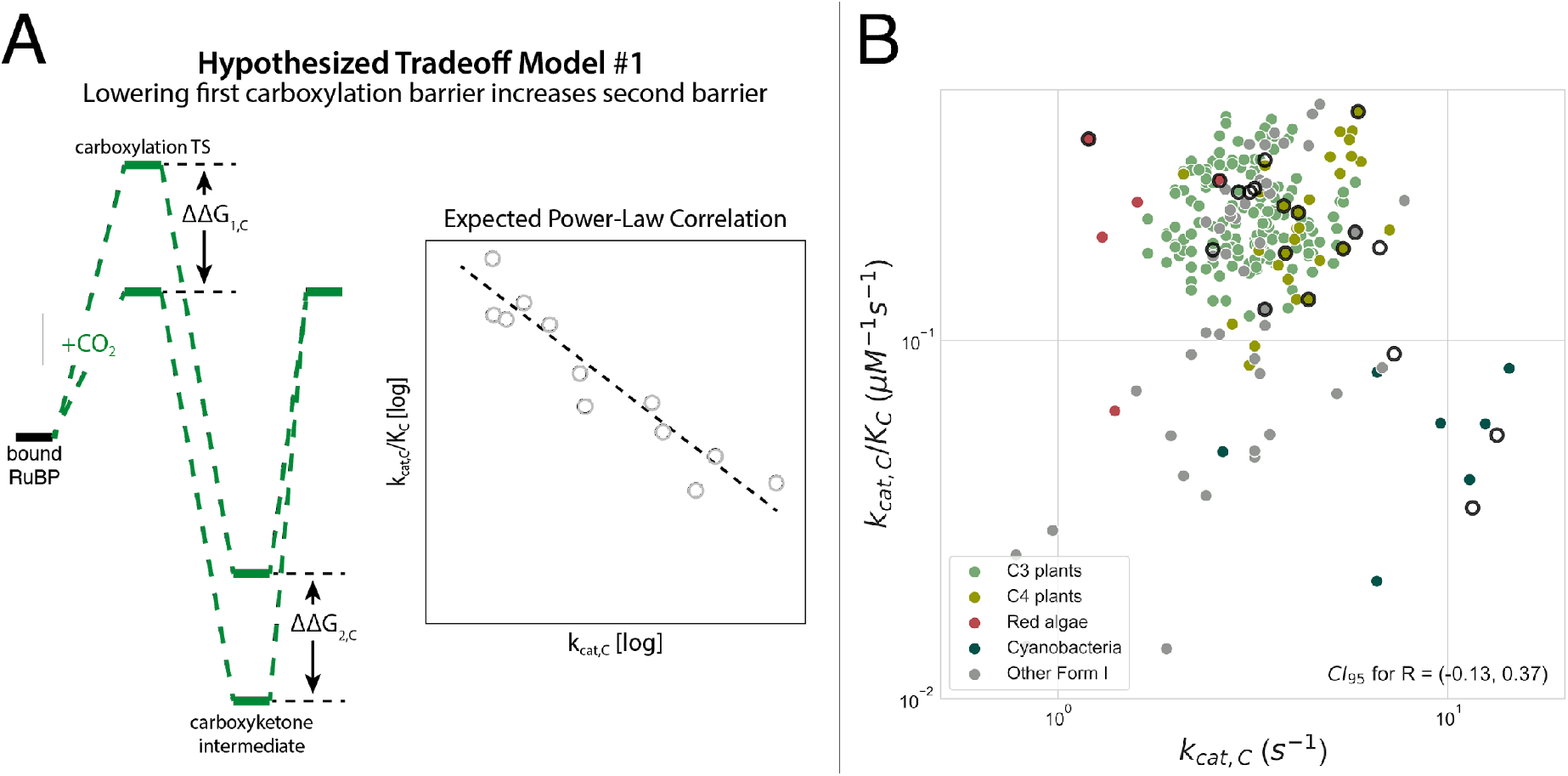
Negative power-law correlation between k_cat,C_ and k_cat,C_/K_C_ is not supported by the extended dataset. In the model diagrammed in panel (A), CO_2_-specific Rubiscos have low barriers to enolization and CO_2_ addition (first effective carboxylation barrier ΔG_1,C_), but lowering the first effective barrier necessarily increases the second effective barrier (ΔG_2,C_), reducing k_cat,C_. In this view, stabilizing the first carboxylation TS also enhances selectivity but also slows carboxylation (Figure S2). ΔG_1,C_ and ΔG_2,C_ should be negatively correlated, which would manifest as negative power-law correlation between k_cat,C_ and k_cat,C_/K_C_ under certain assumptions (SI). (B) The extended dataset does not evidence the expected correlation (R = 0.02, p = 0.8 for Form I enzymes). While previous analyses gave R ≈ −0.9,^6^ the 95% confidence interval for R now includes 0.0. Restricting focus to particular physiologies like C_3_ plants does not recover the expected correlation.

In previous work, k_cat,C_ and k_cat,C_/K_C_ were found to correlate strongly on a log-log scale.^6^ The reported correlation, however, is not strongly supported by our dataset (Figure 6B). The true barrier height to CO_2_ addition depends on the CO_2_ concentration, which could partially explain the apparent lack of correlation. However, correlation is not improved by restricting focus to C_3_ plants for which data is abundant and for which measured leaf CO_2_ concentrations vary by only 20-30% due to variation in CO_2_ conductance and Rubisco activity.^30,31^

Absence of correlation does not necessarily imply the absence of an underlying mechanistic limitation. Rather, if the Rubisco mechanism limits the joint improvement of k_cat,C_ and k_cat,C_/K_C_, much decreased correlation over the extended dataset (R < 0.4) could result from several factors including measurement error, undersampling of Rubiscos with high k_cat,C_ (e.g. from cyanobacteria) or, alternatively, insufficient selection pressure. Diminished correlation, with many points observed below the previous correlation line, suggests that the “mechanistic coupling” model is less likely than “selection within limits” in this case (Figure 2).

The second mechanistic model, wherein faster CO_2_ addition entails faster O_2_ addition,^6^ is well-supported by the extended dataset (Figure 7). This model was previously supported by a power-law correlation between catalytic efficiencies for carboxylation and oxygenation (k_cat,C_/K_C_~(k_cat,O_/K_O_)^2^). As k_cat,C_/K_C_ is exponentially related to the first effective carboxylation barrier (k_cat,C_/K_C_ ~ exp(−ΔG_1,C_)) and k_cat,O_/K_O_ to the first effective oxygenation barrier (k_cat,O_/K_O_ ~ exp(−ΔG_1,O_)), correlation was taken to imply that decreasing the barrier to CO_2_ addition also decreases the barrier to O_2_ addition (Figure 7A). Our dataset supports a similar power-law, albeit with an exponent of ≈1.0 instead of ≈2.0.

**Figure 7:**
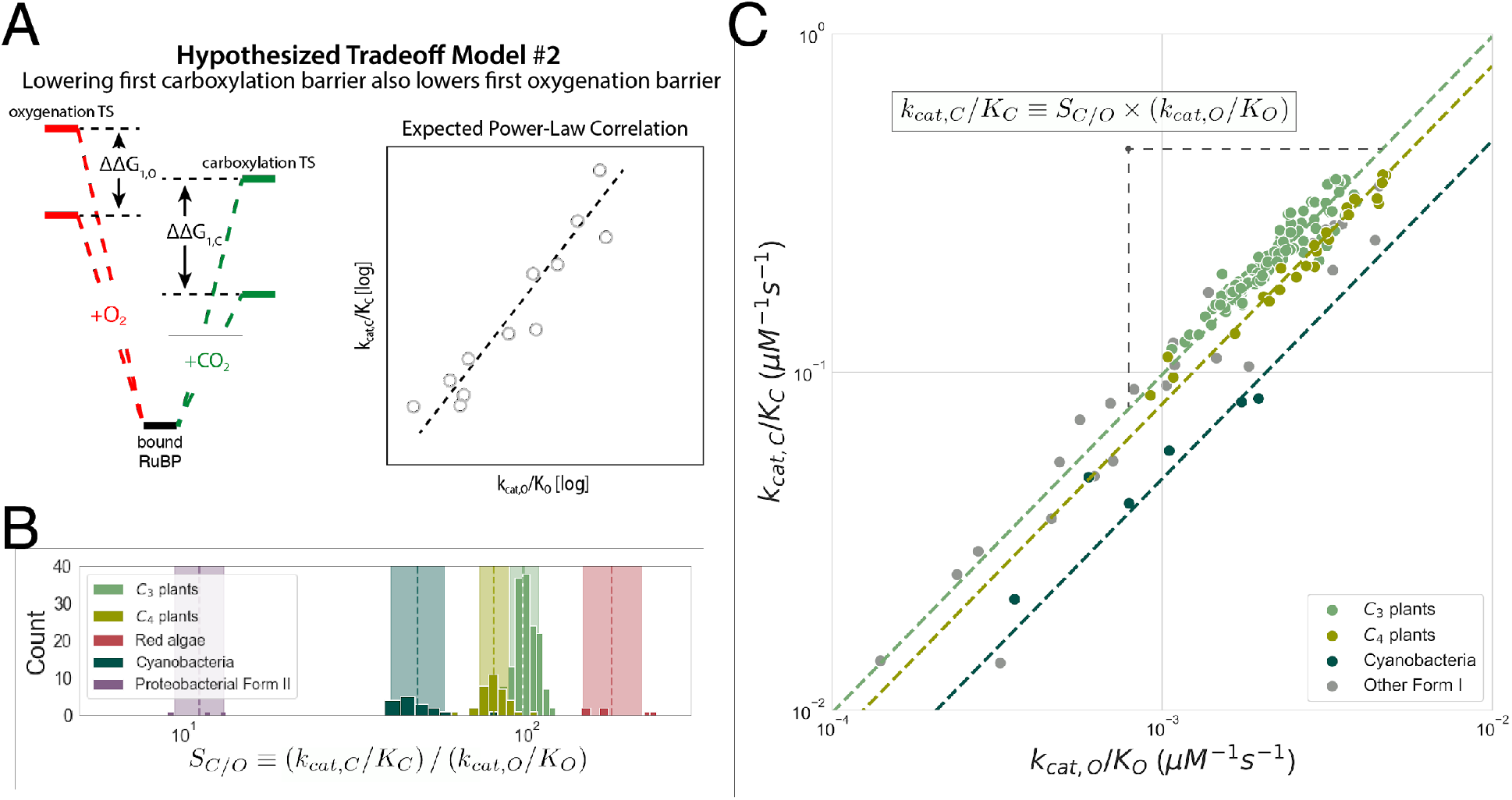
The second mechanistic proposal is remarkably well-supported by the extended dataset. (A) In this proposal, mutations increasing the rate of CO_2_ addition to the Rubisco-RuBP complex also increase the rate of O_2_ addition. In energetic terms, lowering the effective barrier to enolization and CO_2_ addition (ΔG_1,C_) lowers the first effective barrier to O_2_ addition (ΔG_1,O_) as well. Given this model, barrier heights should be positively correlated, which would manifest as a positive linear correlation on a log-log plot of k_cat,C_/K_C_ against k_cat,O_/K_O_. (B) S_C/O_ displays limited variation within physiological groups such as C_3_ and C_4_ plants for which we have substantial data. Dashed lines give the geometric mean of S_C/O_ values. The multiplicative standard deviation, σ*, sets the width of the shaded region. (C) S_C/O_ = (k_cat,C_/K_C_) / (k_cat,O_/K_O_), so restricted S_C/O_ variation implies a power-law relationship (k_cat,C_/K_C_) = S_C/O_ x (k_cat,O_/K_O_). k_cat,C_/K_C_ is strongly correlated with k_cat,O_/K_O_ on a log-log scale (R = 0.94, p < 10). Fitting FI measurements gives k_cat,C_/K_C_ = 119 (k_cat,O_/K_O_)^1.04^. A 95% confidence interval exponent is (0.94, 1.13), which includes 1.0. The geometric mean of measured S_C/O_ values predicts k_cat,O_/K_O_ = (k_cat,C_/K_C_)/S_C/O_ and vice versa. This simple approach accurately predicts of k_cat,O_/K_O_ for FI Rubiscos (prediction R^2^ = 0.80), C_3_ plants (R^2^ = 0.84), C_4_ plants (R^2^ = 0.96) and cyanobacteria (R^2^ = 0.79). Other groups, e.g. red algae, are omitted because of insufficient data.

Again, we found that S_C/O_ varies little among Form I Rubiscos (Figure 3C) and even less within C_3_ plants, cyanobacteria and other physiological groupings (Figures 5A and 7B). S_C/O_ = (k_cat,C_/K_C_) / (k_cat,O_/K_O_) by definition, so the fact that S_C/O_ is approximately constant forces a positive power-law relationship of log(k_cat,C_/K_C_) = log(k_cat,O_/K_O_) + log(S_C/O_). Indeed, Form I enzymes display a remarkably high-confidence power-law relationship between k_cat,C_/K_C_ and k_cat,O_/K_O_ (R = 0.94, *p* < 10^−10^). Since S_C/O_ is the only free parameter in this equation and it is nearly-constant, the geometric mean of S_C/O_ measurements (≈ 90 for Form I Rubiscos) can be used to predict k_cat,O_/K_O_ as (S_C/O_)^−1^ x (k_cat,C_/K_C_). This simple approach, which uses a power-law exponent of 1.0 and prefactor of (S_C/O_)^−1^, predicts Form I k_cat,O_/K_O_ values with R^2^ = 0.80, nearly as accurate as fitting both the prefactor and exponent as free parameters (R^2^ = 0.81). As shown in Figure 7C, predictions of k_cat,O_/K_O_ = (S_C/O_)^−1^ x (k_cat,C_/K_C_) generally improve when considering specific physiological groupings like C_3_ and C_4_ plants because S_C/O_ varies so little within these groups. Assuming roughly constant S_C/O_ forces a 1:1 relationship ΔG_1,C_ = ΔG_1,O_ + C, meaning that decreasing the CO_2_ addition barrier ΔG_1,C_ is associated with an equal decrease to the O_2_ addition barrier ΔG_1,O_.

### Implications for the mechanism of CO_2_/O_2_ discrimination by Rubisco

A 1:1 relationship between effective barriers to CO_2_ and O_2_ addition suggests that a single factor controls both barriers. We offer a simple model based on the mechanism of Rubisco that can produce a 1:1 correlation between barrier heights and constant S_C/O_. In this model, the RuBP-bound active site fluctuates between reactive and unreactive states (Figures 8A). The fraction of enzyme in the reactive state is denoted *ϕ*. In the unreactive state neither oxygenation or carboxylation can proceed. In the reactive state, either gas can react at its intrinsic rate, which does not vary across Rubiscos of the same class (Figure 8B). Since RuBP must undergo enolization in order for carboxylation or oxygenation to occur, *ϕ* may be determined by the degree of enolization of RuBP (SI).

**Figure 8:**
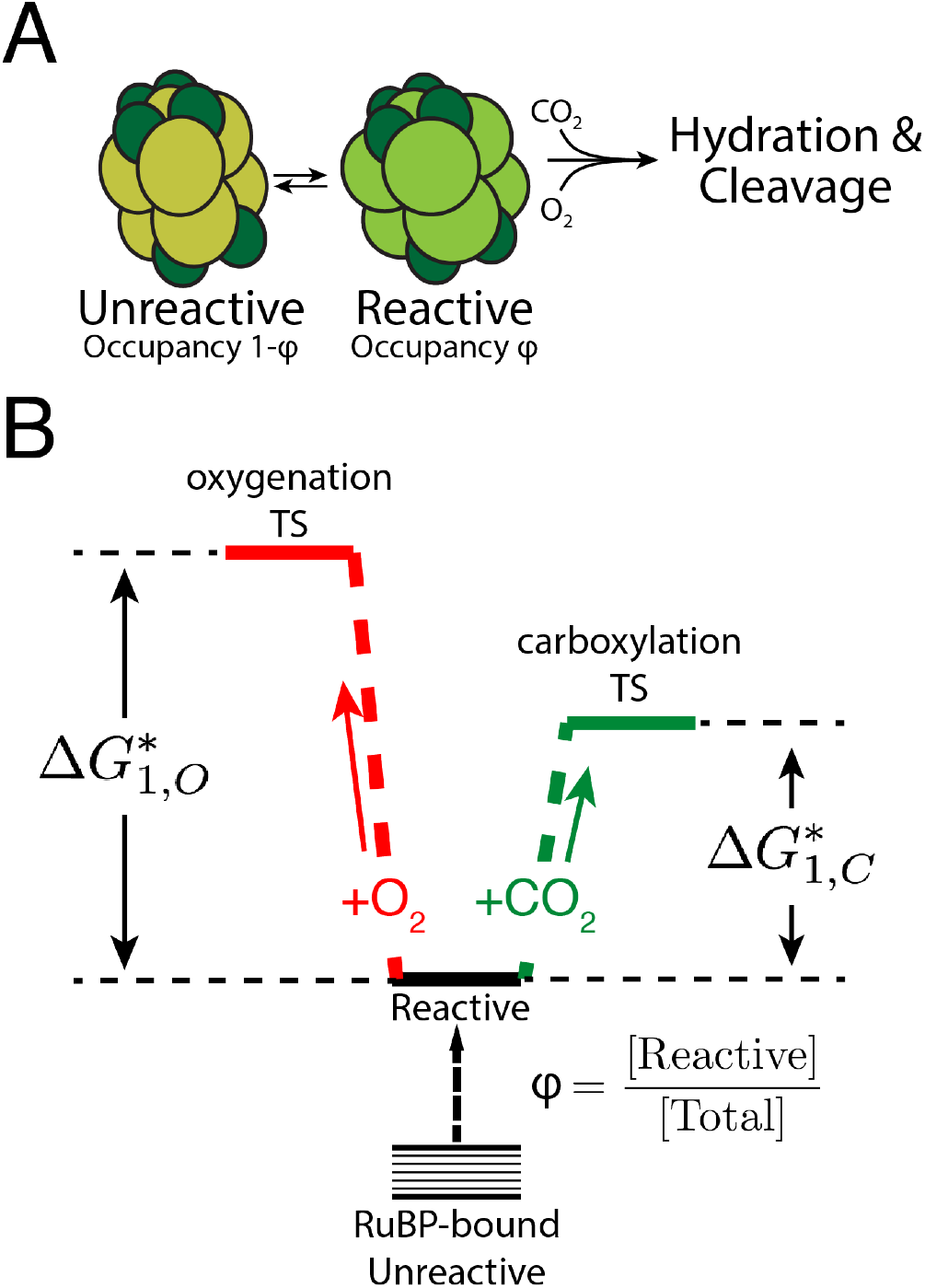
A power-law relationship between k_cat,C_/K_C_ and k_cat,O_/K_O_ can be explained by an active site that fluctuates between “reactive” and “unreactive” states. (A) In this model, CO_2_ and O_2_ react with bound RuBP only when enzyme is in the reactive state, which has occupancy φ. (B) φ can vary between related enzymes. In the reactive state, CO_2_ and O_2_ react with the bound RuBP with intrinsic reactivities ΔG*_1,C_ and ΔG*_1,O_ that do not vary between related Rubiscos. If the difference in intrinsic reactivities (ΔG*_1,O_ − ΔG*_1,C_) is constant, we derive a power-law relationship between k_cat,C_/K_C_ and k_cat,O_/K_O_ with an exponent of 1.0. This relationship requires constant S_C/O_ (SI).

This model can be phrased quantitatively as 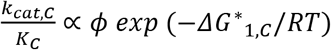 and 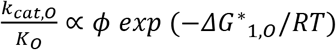 where *ΔG*\*_1,*C*_ and *ΔG*\*_1,*O*_ are the intrinsic reactivities of the RuBP enediolate to CO_2_ and O_2_ respectively. Under this model, S_C/O_ should be roughly constant, which forces a power-law relationship between 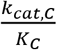 and 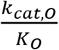 with an exponent 1.0 (Figure 7C, SI). Variation in 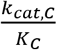 and 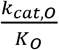 implies that *ϕ* can vary between related Rubiscos, perhaps by evolutionary tuning of the equilibrium constant for RuBP enolization. S_C/O_ is independent of the equilibrium fraction of on-enzyme RuBP enolization, so variation in enolization should affect k_cat,C_/K_C_ and k_cat,O_/K_O_ without altering S_C/O_ (SI). Rather, S_C/O_ is determined by the difference *ΔG*\*_1,*O*_ − *ΔG*\*_1,*C*_ and so changes to the conformation of the RuBP enediolate might explain characteristic differences between S_C/O_ of C_3_ plant and cyanobacterial Rubiscos.^5,8^ See SI for a derivation of this model and further discussion of its implications.

## Discussion

We collected and analyzed literature measurements of ≈300 Rubiscos (Figure 3A). The literature is very phylogenetically-biased, with the readily-purified plant Rubiscos making up >80% of the data (Figure 3B). Despite incomplete coverage, some trends are clear. Rubisco kinetic parameters display extremely limited dynamic range, with multiplicative standard deviations being less than threefold in all cases (Figure 3C). *k_cat,C_* and S_C/O_ appear particularly constrained. Rubisco displays much less *k_cat_* variability than any other enzyme for which sufficient data is available (Figure S5). 97% of *k_cat,C_* values are between 1 and 10 s^−1^ and the highest *k_cat,C_* measured at 25 °C (14 s^−1^, *S. elongatus* PCC 7942^16^) is only ≈20 times the lowest reported Form I value (0.8 s^−1^ from the diatom *Cylindrotheca* N1^32^). Altogether, these data suggest that there is some limitation on the maximum rate of carboxylation by Rubisco in the presence of O_2_.

Focusing on O_2_, we find that measured Rubiscos oxygenate slowly: more than half of *k_cat,O_* measurements are less than 1 s^−1^ and k_cat,C_ is 4 times greater than k_cat,O_ on average (Figures 3C and S4A). Similarly, O_2_ affinity is quite low in general: the median K_O_ is ≈ 470 μM, nearly double the Henry’s law equilibrium of water with a 21% O_2_ atmosphere (≈ 270 μM at 25 °C).

With a multiplicative standard deviation of 1.3, S_C/O_ displays the least variation all Rubisco kinetic parameters (Figure 3C and S4A). Figures 5A and 7B highlight the stereotyped variation in S_C/O_, where C_3_ plant, C_4_ plant, cyanobacterial and red algal enzymes display very limited variation around characteristic S_C/O_ values. All groups have multiplicative standard deviations *σ*\* < 1.25. Nonetheless, Form I Rubiscos are about an order-of-magnitude more CO_2_-specific than the few characterized Form II, III and II/III enzymes (Figure 7B, SI). This might be explained by the prevalence of FII, FIII and FII/III enzymes in bacteria and archaea that fix CO_2_ in anaerobic conditions, where it is doubtful that oxygenation affects organismal fitness. We note, however, that there is substantial variation among measurements of the model FII Rubisco from *R. rubrum* (Figure S4B). This and the paucity of data on non-Form I Rubiscos (Figure 3B) indicates that more measurements are required to evaluate FII, FII/III and FIII enzymes. As such, we focused here on FI Rubiscos, for which data is abundant.

Rubisco kinetics were previously argued to vary in a one-dimensional landscape^6^ and hypothesized to be “nearly perfectly optimized.”^5^ Overall, FI Rubiscos appear less constrained than previously supposed. Figure 4 documents an overall reduction in correlation between FI Rubisco kinetic parameters and the dataset is no longer well-approximated as one-dimensional (Figure S8). Many natural Rubiscos appear suboptimal in plots of k_cat,C_ against S_C/O_ since other enzymes have roughly equal S_C/O_ but higher k_cat,C_ (Figures 5A and S6). Weakened correlations could be due to measurement error and systematic biases, though we find this explanation unlikely because (i) measurements of Form I Rubiscos from similar organisms are broadly consistent (Figure S6); (ii) some correlations remain strong and statistically significant across the entire dataset; (iii) systematic bias towards C_3_ plants would tend to increase correlations; and (iv) standardization of Rubisco assays using stoichiometric inhibitors to quantify active sites should improve data quality over time (SI). Reduced correlations therefore lead us to reject the notion that Rubisco kinetics vary in a strictly one-dimensional landscape and to revisit previous models of mechanistic tradeoff.

The mechanistic models described in Figures 6 and 7 are based on a simple chemical intuition: the intrinsic difficulty of discriminating CO_2_ and O_2_ requires the enzyme to differentiate between carboxylation and oxygenation transition states. The requirement of transition state discrimination is a direct consequence of two assumptions supported by experimental evidence.^22^ Briefly, it is assumed that addition of either gas is irreversible and that there is no binding site for CO_2_ or O_2_ and, thus, no “Michaelis complex” for either gas.^5,6,22,33,34^ If CO_2_ bound a specific site on Rubisco before reacting, it might be possible to modulate K_C_ by mutation without substantially affecting the kinetics of subsequent reaction steps. In the unlikely case that gas addition is substantially reversible,^34,35^ we might expect to find Rubiscos that evolved enhanced selectivity by energy-coupled kinetic proofreading. Energy coupling can enable amplification of selectivity due to differential CO_2_ and O_2_ off-rates.^36^ The fact that no such Rubiscos have been found suggests that gas addition is irreversible or that CO_2_ and O_2_ off-rates are incompatible with kinetic proofreading in some other way.^6,37^

As Rubisco likely does not bind CO_2_ directly, it was hypothesized that high CO_2_-specificity (large S_C/O_) is realized by discriminating between the first carboxylation and oxygenation transition states, i.e. between the developing carboxyketone and peroxyketone (Figures S1 and S2).^5^ A late carboxylation transition state would be maximally discriminable because the developing carboxylic acid is distinguishable from the peroxyl group of the oxygenation intermediate. The extraordinarily tight binding of the carboxyketone analog CABP to plant Rubisco provides strong support for a late carboxylation transition state.^5^ Since a late transition state resembles the carboxyketone intermediate, it was argued that CO_2_-specific Rubiscos must tightly bind the intermediate, slowing the subsequent reaction steps and restricting k_cat,C_ (Figure S2).^5^

As k_cat,C_/K_C_ is related to the effective barrier to enolization and CO_2_ addition (ΔG_1,C_) and k_cat,C_ is related to the effective barrier to hydration and cleavage (ΔG_2,C_, Figure 1D), an energetic framing of this model argues that lowering ΔG_1,C_ (increasing k_cat,C_/K_C_) entails increasing ΔG_2,C_ (lowering k_cat,C_, Figure 6A).^6^ Despite nuanced differences, we collectively term these models k_cat,C_-K_C_ coupling due to the hypothesized coupling of carboxylation kinetics. Though these models are motivated by the need to discriminate between CO_2_ and O_2_, they invoke a tradeoff between carboxylation steps only. That is: specificity requires tighter binding of the carboxylation intermediate, which slows downstream processing of that same intermediate, irrespective of O_2_.

Three correlations previously supported k_cat,C_-K_C_ coupling: correlation between k_cat,C_ and S_C/O_, between k_cat,C_ and K_C_, and between k_cat,C_ and k_cat,C_/K_C_. k_cat,C_ and S_C/O_ remain negatively correlated in our larger dataset, but more weakly than previously observed (Figure 5A). The same is true for k_cat,C_ and K_C_ (Figure 5B) and for k_cat,C_-k_cat,C_/K_C_ (Figure 6B). Rather than arguing for strong coupling of carboxylation kinetics, Figure 5 highlights the stereotyped variation in S_C/O_ described above. We interpret weakened correlations as implying that carboxylation kinetics are not strictly coupled. Considering residuals of the k_cat,C_-K_C_ fit (Figure S9) shows that outliers include recent measurements of cyanobacterial^38^ and diatom^39^ Rubiscos, which fall well-below the fit line. This is consistent with a “selection within limits” view of k_cat,C_-K_C_ coupling (Figure 2B).

The second mechanistic tradeoff model posits that faster CO_2_ addition to the Rubisco-RuBP complex necessarily allows faster O_2_ addition. This model was previously supported by a positive power-law correlation between the catalytic efficiencies for carboxylation and oxygenation (k_cat,C_/K_C_ and k_cat,O_/K_O_ respectively), which can be understood as a positive coupling of the effective barriers to enolization and gas addition for CO_2_ and O_2_ (ΔG_1,C_ and ΔG_1,O_, Figure 7A). We showed that extremely limited and stereotyped variation in S_C/O_ = (k_cat,C_/K_C_) / (k_cat,O_/K_O_) necessitates a power-law correlation with exponent 1.0 between k_cat,C_/K_C_ and k_cat,O_/K_O_ (Figure 7B-C). An exponent of 1.0 implies that decreasing ΔG_1,C_ (enabling faster carboxylation) requires a *roughly equal* reduction to ΔG_1,O_ (enabling faster oxygenation as well). Although several research groups have attempted to isolate improved Rubisco mutants, none of the mutants examined so far exceed wild-type enzymes on these axes (Figure S11).

A power law relation with an exponent of 1.0 can be seen as resulting from an active site that fluctuates between a reactive and unreactive state (Figure 8A). This coarse-grained model is motivated by the Rubisco mechanism in two ways. Since Rubisco likely does not bind CO_2_ or O_2_ directly, active site concentrations are determined by solution concentrations (e.g. in the chloroplast stroma). Rubisco could close the active site to diffusion to limit O_2_ entry,^40^ but this would slow carboxylation as well. Similarly, RuBP must enolize for oxygenation or carboxylation to proceed (Figure 1A) so modulating the degree of enolization would affect both reaction pathways equally.^14,15^ In either case, the average occupancy of the reactive state mechanistically couples the rates of CO_2_ and O_2_ addition (Figure 2A) and throttles the subsequent steps of carboxylation and oxygenation equally (Figure 8).

In previous work, where Rubisco kinetics were thought to vary in a one-dimensional landscape, setting k_cat,C_ determined all other kinetic parameters.^6^ In this setting, it was argued that Rubisco kinetic parameters are determined by the prevailing CO_2_ and O_2_ concentrations since a unique choice of parameters on the one-dimensional curve maximizes the net rate of carboxylation.^6^ Since the data is no longer clearly one-dimensional, we cannot argue that Rubisco is “perfectly optimized” to match prevailing concentrations. Moreover, the model presented in Figure 8 sets no upper limit on k_cat,C_, suggesting that selection for increased carboxylation in the absence of O_2_ could produce Rubisco mutants with superlative k_cat,C_ values (i.e. k_cat,C_ ≫ 15 s^−1^). Such enzymes might be found in anaerobic bacteria and would be of interest in probing the limits of Rubisco catalysis.

The prospect of engineering an improved Rubisco is tantalizing, not only because it could plausibly improve crop yields,^18^ but also because the task tests our understanding of enzymes on a very basic level. It is clear from the data presented here that there is some evolutionary constraint on Rubisco catalysis. Surely a superlative Rubisco would have arisen if it was mutationally accessible from existing enzymes. More detailed biochemical investigation of naturally-occuring Rubiscos will help delineate the evolutionary constraints imposed on Rubisco kinetics. Still, the Rubisco large subunit displays extremely limited sequence variation.^41^ Perhaps exploring a wider swath of sequence space via protein engineering techniques^42–44^ would enable strict improvements to Rubisco kinetics? We argue that biochemical and bioengineering techniques should be used in concert to probe the limits of Rubisco catalysis and propose several avenues of future research to evaluate the prospects of Rubisco engineering.

First, the kinetics of non-plant Rubiscos should be characterized more thoroughly. These should include the Form II, III and II/III enzymes of bacteria and archaea as well as FI enzymes of bacteria and diverse eukaryotic autotrophs.^13,39^ Ideally these enzymes would be chosen in a manner that maximizes sequence and phylogenetic diversity^45^ and characterized for their binding (e.g. of RuBP and CABP) and catalytic activity (measuring k_cat,C_, K_C_, k_cat,O_, K_O_ and S_C/O_) as a function of temperature and pH.^29,46,47^ A facile assay for direct measurement of oxygenation would also reduce the number of assumptions made in measuring and analyzing Rubisco kinetics.^29^ These data would help resolve whether Rubisco isoforms display characteristic differences in catalytic potential. It is possible, for example, that non-Form I enzymes are subject to different constraints than FI Rubiscos and might serve as useful chassis for engineering.

It is also important to revisit the classic experiments undergirding our understanding of the Rubisco catalytic mechanism, especially those supporting the central assumptions that (i) there is no Michaelis complex for CO_2_ or O_2_ and (ii) that gas addition is irreversible.^22,34,35^ These assumptions substantially constrain CO_2_ specificity. If we were to find Rubiscos for which these assumptions are relaxed, they might serve as a basis for engineering a fast-and-selective carboxylase. On the other hand, all Rubiscos may share the same limitations. Since these limitations are likely described as couplings between transition state barriers (as in Figures 7 and 8) measurements of barriers heights for a wide variety of Rubiscos would enable more direct testing of tradeoff models. One avenue for drawing inference about barrier heights is by measuring the binding energies of intermediate and transition state analogs.^5,48^ Kinetic isotope effects for CO_2_ and O_2_ report indirectly on the relevant barriers^49^ and can be measured by mass spectrometry.^50^ Investigating the relationship between transition state barriers and kinetic parameters will help delineate which reaction steps limit carboxylation and oxygenation in different Rubisco lineages.^5^

There remains some disagreement about the precise ordering of carboxylation steps^5,14,15^ and the mechanism of oxygenation is not well understood.^48^ Chemical reasoning about the mechanisms of Rubisco carboxylation and oxygenation would benefit from progress in structural biology - intermediate and transition state analogs should be used to capture the active site at various points along the reaction trajectory. ^14,40,48,51^ If experiments and structural analyses confirm that the above assumptions hold for all Rubiscos, it would greatly limit our capacity to engineer Rubisco and strongly suggest that alternative strategies for improving carbon fixation should be pursued.^19,52–54^ If, however, these assumptions are invalidated, many enzyme engineering strategies would become viable. Such data and analyses will be instrumental in guiding the engineering of carbon fixation for the next decade.

## Supporting information

Dataset S2 Merged Dataset

Dataset S3 BRENDA Kinetic Data

Dataset S4 Full Merged Dataset with Mutants

## Acknowledgements

We would like to thank Uri Alon, Kapil Amarnath, Doug Banda, Arren Bar-Even, Cecilia Blikstad, Jack Desmarais, Woodward Fischer, Vahe Galstyan, Laura Helen Gunn, Itai Halevy, Robert Nichols, Elad Noor, Jeremy Roop, Yonatan Savir, Patrick Shih, Daniel Stolper, Dan Tawfik, Guillaume Tcherkez, Tsvi Tlusty and Renee Wang for helpful conversations and comments on the manuscript. This research was supported by the US National Science Foundation (grant MCB-1818377); the European Research Council (project NOVCARBFIX 646827); the Israel Science Foundation (grant No. 740/16); the ISF-NRF Singapore joint research program (grant No. 7662712); Beck-Canadian Center for Alternative Energy Research; Dana and Yossie Hollander; Ullmann Family Foundation; Helmsley Charitable Foundation; The Larson Charitable Foundation; Wolfson Family Charitable Trust; Charles Rothschild; Selmo Nussenbaum. R.M. is the Charles and Louise Gartner professional chair. A.I.F. was supported by a National Science Foundation Graduate Research Fellowship. Y.M.B. is an Azrieli fellow.

