## Supplemental text for "Revisiting tradeoffs between Rubisco kinetic parameters"

**Supplementary Information: Revisiting tradeoffs in Rubisco kinetic parameters** 1
- Previous Literature on Tradeoffs in Rubisco Catalysis 3
  - Rubisco Folding, Activation, and Catalytic Limitations 3
  - Measurement of Rubisco Kinetic Parameters 4
  - Previous Literature on Tradeoffs in Rubisco Catalysis 5
- Description and Analysis of the Extended Dataset 7
  - Collection and Curation of the Extended Dataset 7
  - Restricted Variation in Rubisco Kinetic Parameters 8
  - Correlation and Regression Analyses 9
  - Use of Log-Scale Correlations 9
  - Principal Component Analysis 11
  - Measurement Error and Systematic Error in Rubisco Measurements 11
  - Correlations with kcat,O 12
- Relationship Between Tradeoff Models and the Microscopic Kinetics of Rubisco 13
  - Derivation of Rubisco Kinetic Equations 13
  - Catalytic Efficiencies (kcat/KM) are Related to the First Effective Barrier to Carboxylation and Oxygenation 13
  - kcat,C is Related to the Second Effective Barrier for Carboxylation 14
  - Interpretation of Power-Law Correlation Between kcat,C/KC and kcat,O/KO 15
  - Derivation of the Active Site Gating Model 15
  - Implications of the Active Site Gating Model 16
  - Characterized Rubisco Mutants Do Not Exceed Wild-Type Tradeoffs 17
- Supplementary Tables 18
- Supplementary Figures 20

References 31

### Previous Literature on Tradeoffs in Rubisco Catalysis

#### Rubisco Folding, Activation, and Catalytic Limitations

Rubisco is likely the most abundant enzyme in nature, being the central enzyme of the Calvin-Benson-Bassham cycle responsible for nearly all annual carbon fixation.[1,2] Rubisco is a notoriously complex enzyme that depends on multiple chaperones for folding, assembly and catalysis,[3–6] requires a post-translational covalent modification to become active, and is inhibited by multiple metabolites, including its own five-carbon substrate.[5–7] Activation consists of carbamylation of an active-site lysine and binding of a catalytic $Mg^{2+}$. If the five-carbon substrate ribulose 1,5,-bisphosphate (RuBP) binds prior to activation, the enzyme cannot become activated. Diverse ATP-coupled chaperones, collectively termed Rubisco activases, have evolved to eject RuBP from the active site and permit activation.[5,6]

Once activated, all known Rubiscos catalyze carboxylation and oxygenation of RuBP through a multistep mechanism (Figure S1). Both carboxylation and oxygenation of RuBP are energetically favorable,[8] but only carboxylation is considered productive because it incorporates carbon from $CO_2$ into precursors that can generate biomass. Oxygenation is often portrayed as counterproductive as it occupies Rubisco active sites and yields a product (2-phosphoglycolate, 2PG) that is not part of the CBB cycle and must be recycled through metabolically-expensive photorespiration at a loss of carbon.[9–11] As photorespiration is known to play a role in signalling and nitrogen metabolism in plants,[10–12] there is disagreement about whether oxygenation and photorespiration are "wasteful." However, many phototrophs have evolved $CO_2$ concentrating mechanisms (CCMs) that suppress oxygenation by elevating the $CO_2$ concentration near the Rubisco active site.[9,13,14] Moreover, efforts to improve photorespiration through pathway engineering have proved fairly successful in cyanobacteria and plants,[15–18] implying that the products of oxygenation by Rubisco do in fact impinge on autotrophic growth, albeit in a complex manner. Despite the fact that many autotrophs depend on Rubisco carboxylation for growth, all known Rubiscos are relatively slow carboxylases that fail to exclude $O_2$ (Figures 1A-B and S1).

While it is often claimed that Rubisco is "slow" and "non-specific" it is actually the case that Rubisco is a slightly below-average enzyme in terms of carboxylation $k_{cat,C}$ and roughly average in terms of $k_{cat}/K_M$.[19,20] Moreover, Rubisco catalysis is calculated to enhance the rate of RuBP carboxylation by $\approx 10^{16}$ fold. A 15-16 order-of-magnitude rate enhancement is quite impressive, especially in comparison to canonical "perfect" enzymes like carbonic anhydrase, superoxide dismutase and triose-phosphate isomerase that display rate enhancements on the order of $10^7$-$10^9$ fold.[20]

Rubisco is not "slow" in an absolute sense. Rather, we are surprised that Rubisco is not faster given its centrality to life and its abundance.[19,20] Rubisco is routinely measured to comprise upwards of 20% of total soluble leaf protein in $C_3$ plants.[21] Moreover, oxygenation by Rubisco competitively inhibits carboxylation[9,22,23] and produces a product, 2PG, that is metabolized by decarboxylation.[11] Therefore, the abundance of $O_2$ in present-day atmosphere (21%)

suppresses the net rate of carboxylation by Rubisco. These conditions might be expected to select for a Rubisco that is both fast-carboxylating (having high $k_{cat,C}$) and highly specific (also having superlative $S_{C/O}$), but such an enzyme has not yet been found. Rather, $CO_2$ concentrating mechanisms elide these problems by elevating the $CO_2$ concentration near the Rubisco active site, which promotes fast carboxylation and competitively inhibits oxygenation.[9,14]

The curious slowness and non-specificity of such an abundant and central enzyme, combined with the fact that CCMs evolved multiple times in diverse lineages,[14] suggests that perhaps Rubisco itself cannot be strictly improved - i.e. that some physicochemical limitation imposes a tradeoff on the enzyme such that it cannot be both fast and specific.[22,24–26] As discussed in the main text and below, a negative correlation between the maximum carboxylation rate, $k_{cat,C}$, and $CO_2$-specificity, $S_{C/O}$, is routinely cited as evidence for this view. Two broad families of mechanistic tradeoff models have been formulated to explain apparent tradeoffs in Rubisco catalysis. All of these analyses are fundamentally based on correlations between measured Rubisco kinetic parameters. We collected and examined a large dataset containing kinetic data for ≈300 Rubiscos to examine whether these mechanistic models are consistent with new data. In the foregoing sections we describe

#### Measurement of Rubisco Kinetic Parameters

Here we use $k_{cat,C}$ and $k_{cat,O}$ to denote the maximum catalytic rates (measured in units of $s^{-1}$) for carboxylation and oxygenation respectively. $K_C$ and $K_O$ denote the effective Michaelis constants (half-saturation concentrations in μM units) for carboxylation and oxygenation. The specificity factor $S_{C/O} = (k_{cat,C}/K_C) / (k_{cat,O}/K_O)$ is a unitless measure of the relative preference for $CO_2$ over $O_2$ (Figure 1A). It should be noted the relationship between $S_{C/O}$ and the per-active site rates of carboxylation ($R_C$) and oxygenation ($R_O$) is complex and nonlinear, as discussed below. Readers are cautioned against the false inference that higher $S_{C/O}$ is “better.” The optimal $S_{C/O}$ value depends strongly on the $CO_2$ and $O_2$ concentrations as well as the absolute values of $k_{cat,C}$, $k_{cat,O}$, $K_C$, and $K_O$.

The per-active site rates of carboxylation ($R_C$) and oxygenation ($R_O$) are calculated as:

$$R_C = k_{cat,C}(1 + \frac{K_C}{[CO_2]} + \frac{K_C/[CO_2]}{K_O/[O_2]})^{-1}$$

$$R_O = k_{cat,O}(1 + \frac{K_O}{[O_2]} + \frac{K_O/[O_2]}{K_C/[CO_2]})^{-1}$$

These equations take the form of an irreversible Michaelis-Menten type relationship with mutual competitive inhibition. That is, both carboxylation and oxygenation are irreversible - both reactions are associated with strongly negative $\Delta_r G'^m$ values[8] - and both reactions mutually inhibit each other competitively by occupying the same active site.[23,27,28] The rate laws above are independent of the RuBP concentration, implicitly assuming that RuBP is present in saturating concentrations. When RuBP is saturating, per-enzyme rates can be multiplied by the concentration of active Rubisco ([E]) to calculate the total rates of carboxylation $V_C$ and $V_O$.

Rubisco assays are challenging to perform, not least because both inorganic substrates are gasses. Moreover, due to the activation step, the number of active sites is not reliably quantified by measuring protein concentration. A stoichiometric inhibitor, 2-carboxyarabinitol 1,5-bisphosphate (CABP), is typically used in contemporary assays to quantify the number of active sites after activation in the presence of $CO_2$ and $Mg^{2+}$.[29] The carboxylation rate of activated Rubisco can then be measured by a variety of methods, e.g. as incorporation of $^{14}CO_2$ into acid-stable material over time.[30] Performing these assays in sealed vessels at a variety of $CO_2$ and $O_2$ partial pressures enables fitting of $k_{cat,C}$, $K_C$, and $K_O$ on the assumption that the Michaelis constant for $O_2$ ($K_O$) is equal to the half-maximum inhibitory $O_2$ concentration.

The median $K_O$ value is ≈ 470 μM (Figure 3C), nearly double the ≈ 270 μM Henry's law equilibrium of water with 21% $O_2$ atmosphere at 25 °C.[31] As such it can be difficult to saturate Rubisco with $O_2$ in order to measure $k_{cat,O}$. Moreover, the assay described above measures only the incorporation of $CO_2$ (not $O_2$) and so a different assay is required to infer $k_{cat,O}$. This achieved by an independent measurement of $S_{C/O}$, after which $k_{cat,O}$ is calculated as ($k_{cat,C}/K_C$) / ($S_{C/O}/K_O$). $S_{C/O}$ assays take advantage of the fact that $\frac{R_C}{R_O} = S_{C/O} \times \frac{[CO_2]}{[O_2]}$in the limit of low $CO_2$ and $O_2$ concentrations (see below). To measure $S_{C/O}$, the rate of $O_2$ and $CO_2$ incorporation are measured simultaneously in the same sealed assay vessel.[32,33] Plotting $R_C/R_O$ against $[CO_2]/[O_2]$ gives $S_{C/O}$ as the slope of linear fit passing through $R_C/R_O = 0$.

#### Previous Literature on Tradeoffs in Rubisco Catalysis

Correlation between $S_{C/O}$ and other Rubisco kinetic parameters is often cited to motivate the notion that tradeoffs mediated by the catalytic mechanism strictly constrain Rubisco's catalytic potential.[22,24–26,34,35] Tcherkez et al. 2006 focuses primarily on the correlation between $S_{C/O}$ and $k_{cat,C}$ and argues that this relationship is best-explained by the intrinsic difficulty of discriminating between $CO_2$ and $O_2$, both of which are small, volatile and similarly nonpolar molecules.[25] As a result, Tcherkez et al. argue, discrimination must occur between the carboxylation and oxygenation transition states (TS). An "advanced, product-like" carboxylation TS allows for maximum discrimination because the developing carboxylate intermediate is more readily distinguished from the peroxyacid of the oxygenation intermediate. However, tight-binding of a product-like intermediate is expected to limit the throughput of the subsequent, rate-limiting, reaction steps by slowing the release of the carboxyketone carboxylation intermediate (Figure S2).[25]

This explanation is consistent with a number of lines of evidence given by Tcherkez et al., including (i) correlation between $S_{C/O}$ and $k_{cat,C}$, (ii) the measured temperature dependence of Rubisco carboxylation and oxygenation, (iii) trends in the carbon kinetic isotope effect and (iv) the extremely tight binding of the carboxyketone analog CABP.[25] If there is a tradeoff between carboxylation rate and specificity, Rubisco may be "nearly perfectly optimized" to suit environmental $CO_2$ and $O_2$ concentrations and maximize the rate of carbon gain in each host organism. However, data is limited for all of these claims and linear correlation plots of Tcherkez et al. 2006 have notable outliers in the cyanobacterial Rubiscos, which are faster than the overall trend would predict.[25]

Savir et al. 2010 - a study in which the last author of this work participated - can be understood as starting from a simple motivating question: why is there so little variation in Rubisco kinetics? The extremely limited variation in Rubisco kinetics is highlighted by Figure 3C, where multiplicative standard deviations for all measured kinetic parameters are less than threefold (among Form I Rubiscos). If all kinetic parameters ($S_{C/O}$, $K_C$, $k_{cat,C}$, $K_O$ and $k_{cat,O}$) were free variables, then they would vary in a 4-dimensional space (not 5-dimensional because $S_{C/O}$ is wholly determined by the other parameters). Savir et al. 2010 uses PCA in the space of log-transformed kinetic parameters to show that the kinetics of ≈20 diverse Rubiscos are roughly one dimensional, meaning that there is only one free variable in the system. In other words, $S_{C/O}$, $K_C$, $K_O$ and $k_{cat,O}$ can be calculated with high accuracy from $k_{cat,C}$ alone.

Savir et al. further argue that the kinetics associated with particular Rubiscos, e.g. those from $C_3$ plants or cyanobacteria, can be understood as maximizing the net rate of carboxylation $f = R_C - 0.5\ R_O$ in that host. Concretely: cyanobacteria possess a CCM that is thought to produce [$CO_2$] > 250 μM,[9] while $C_3$ plants have no CCM and so $C_3$ plant Rubiscos experience ambient $CO_2$ and $O_2$ concentrations. Since their small dataset is roughly one-dimensional, there is a unique choice of kinetic parameters that maximizes $f$ and, conversely, an optimal $CO_2$ concentration that maximizes $f$ for each set of kinetic parameters. Based on their analysis, $C_3$ plant Rubiscos appear optimized for ambient concentrations, while Rubiscos from cyanobacteria and other CCM-bearing organisms appear to be optimized for elevated $CO_2$ concentrations.

Based on the established kinetic model of Rubisco activity and some assumptions, Savir et al. 2010 note that the catalytic efficiencies for carboxylation and oxygenation are related to "effective barriers" to $CO_2$ and $O_2$ addition ($k_{cat,C}/K_C \sim \exp(-\Delta G_{1,C})$; $k_{cat,O}/K_O \sim \exp(-\Delta G_{1,O})$).[22] Similarly, given assumptions described below, $k_{cat,C}$ is related to the height of a second "effective barrier" that represents hydration and cleavage of the bound carboxylation intermediate ($k_{cat,C} \sim \exp(-\Delta G_{2,C})$, ERC in Figure S1). These relationships are diagrammed in Figure 1D and derived below.

Savir et al. show that $k_{cat,C}$ correlates negatively with $k_{cat,C}/K_C$ on a log-log plot and propose the following explanation. $k_{cat,C}/K_C$ and $k_{cat,C}$ are exponentially related $\Delta G_{1,C}$ and $\Delta G_{2,C}$ respectively.[22] Power-law correlation between these parameters implies that $\Delta G_{1,C}$ and $\Delta G_{2,C}$ sum to a constant. This was taken to mean that a maximum "deformation energy" that can be applied to RuBP and this energy must be partitioned between two reaction steps. This proposed tradeoff $k_{cat,C}/K_C$ and $k_{cat,C}$ is similar to the proposal of Tcherkez et al. and could produce the observed correlation between $S_{C/O}$ and $k_{cat,C}$ without reference to the kinetics of oxygenation.[22] We therefore refer to these two proposals collectively as $k_{cat,C}$-$K_C$ coupling. Savir et al. also show that $k_{cat,C}/K_C$ is positively correlated with $k_{cat,O}/K_O$ on a log-log plot, implying that there is some coupling between $\Delta G_{1,C}$ and $\Delta G_{1,O}$ (Figures 1B and 7A). This correlation implies that mutations increasing the rate of $CO_2$ addition to the ERC complex (Figure S1) also increase the rate of $O_2$ addition.

Savir et al. note that their dataset is too small to discriminate between “mechanistic coupling” or “selection within limits” models diagrammed in Figure 2, but argue that “selection within limits” is more likely because kinetic parameters that affect the net rate of carboxylation tend to correlate more strongly than other pairs. For example, $K_C$ and $K_O$ only affect the net carboxylation rate through their ratio ($K_C/K_O$) and correlate much less than $k_{cat,C}$ and $K_C$ in their dataset. In further support of this view, Savir et al. also show maximum per active site oxygenation rate $k_{cat,O}$ correlates poorly with the other parameters, suggesting that selection is required for correlation (“selection within limits” model, Figure 2B) because carboxylation is certainly the primary trait selected for in autotrophs.

Altogether, Tcherkez et al. 2006 and Savir et al. 2010 put forward two distinct families of models that could explain why a superlatively fast-and-selective Rubisco has not yet been found. In the first family, which we term $k_{cat,C}$-$K_C$ coupling, the steps of the carboxylation mechanism are negatively coupled such that increasing carboxylation $k_{cat,C}$ forces an increase in $K_C$ (i.e. reduced $CO_2$-affinity) and, therefore, a decrease in $S_{C/O}$. These models are motivated by the need for $CO_2/O_2$ discrimination, but are strictly independent of oxygenation kinetics. That is, no changes to oxygenation kinetics are invoked and no correlation with oxygenation kinetics is necessarily predicted. The second family of models, in contrast, invokes a coupling between the rates of $CO_2$ and $O_2$ addition to the Rubisco-RuBP complex. In this model, increasing the rate of $CO_2$ addition ($k_{cat,C}/K_C$) forces an increase to the $O_2$ addition rate ($k_{cat,O}/K_O$).

Both of these models appeal to physico-chemical intuition. However, their mechanistic claims are far from straightforward and warrant analysis in the light of new data. More than 250 Rubisco variants have been characterized since 2010. Here we examine whether new data supports or contradicts the models collectively advanced by Tcherkez et al. 2006 and Savir et al. 2010.[22,25]

### Description and Analysis of the Extended Dataset

#### Collection and Curation of the Extended Dataset

The raw extended dataset including kinetic parameter values from the primary literature as well as temperature and pH of measurements and manual annotations is given as Dataset S1. Experimental error was recorded when reported. We attempted to exhaustively extract kinetic parameters for wild-type Rubiscos. Several values for mutant and hybrid enzymes were also extracted in the process. We reviewed the primary literature and manually annotated cases where the underlying data did not appear trustworthy, e.g. measurements taken before the active site stoichiometry of Form I Rubiscos was understood.[36] Equivocal measurements, mutant enzymes and some values measured at temperatures other than 25 $^oC$ were filtered before further analysis.

In cases where experimental error was not reported, we assumed that measurement error scales with the measured value. Figure S3 supports this assumption, plotting reported error against the measured value for all commonly-measured Rubisco kinetic parameters. We calculated the mean coefficient of variation (CV, the ratio of the standard deviation to the mean) for each commonly measured kinetic parameter ($S_{C/O}$, $k_{cat,C}$, $K_C$ and $K_O$) and assumed this CV to assign a standard deviation in cases of unreported error.

In some cases multiple values for the same parameter were found in the same reference, for example multiple similar $S_{C/O}$ values (and standard deviations) for *R. rubrum* and *S. oleracea* Rubiscos.[37] When these values were similar, they were combined into a single mean and standard deviation by bootstrapping. It was assumed that measurements were normally distributed with the reported standard deviation and means. $10^4$ values were randomly sampled from the implied distribution (i.e. one per measurement being combined) and a posterior mean and standard deviation was calculated.

$S_{C/O}$ is often measured at pH 8.3 while kinetic parameters $k_{cat,C}$, $K_C$ and $K_O$ are usually measured near pH 8.0.[38,39] It is commonly assumed that $S_{C/O}$ is pH-independent after Jordan et al.[37] We made the same assumption and combined kinetic measurements and $S_{C/O}$ values from the same reference. This assumption is used to calculate $k_{cat,O}$ from the other four parameters as $k_{cat,O} = (k_{cat,C}/K_C) / (S_{C/O}/K_O)$. Given the combined data, we calculated a 95% confidence interval on $k_{cat,O}$ by $10^4$-fold bootstrapping (Methods) in cases where $S_{C/O}$, $k_{cat,C}$, $K_C$ and $K_O$ were measured in the same reference. When sufficient data was available we also calculated the 95% confidence intervals for $k_{cat,C}/K_C$ and $k_{cat,O}/K_O$ by bootstrapping. The median of bootstrapped values was used to represent $k_{cat,O}$, $k_{cat,C}/K_C$ and $k_{cat,O}/K_O$ in plots (e.g. in Figures 5-7). Inferred values were checked for internal consistency and consistency with literature values before analyses. We refer to the resulting dataset - where data from individual references are merged and unmeasured values are inferred - as the extended dataset.

The complete extended dataset is given in Dataset S2 and source code used for dataset normalization and error inference is available at https://github.com/flamholz/rubisco. Dataset S2 is filtered such that it contains only measurements of native (wild-type, non-mutant) enzymes taken near pH 8 and at 25 °C. An unfiltered version of Dataset S2 is given as Dataset S4. This repository contains data about mutant and hybrid Rubiscos as well as some values measured away from 25 °C. We did not search comprehensively for measurements of mutants or measurements at non-standard temperatures, so Dataset S4 is likely far from complete in this respect. Supplementary datasets are described in Tables S1 and S2.

#### Restricted Variation in Rubisco Kinetic Parameters

The extended dataset displays little variation among all carboxylation and oxygenation parameters, with central values varying over less than fourfold in all cases (Figure S4). For FI Rubiscos, the multiplicative standard deviation ($\sigma^*$) is less than 3.0 for all measured kinetic parameters (Figures 3C and S4). Rubisco differs notably from other enzymes in this regard: for most reaction classes, kinetic parameters vary over 2-3 orders of magnitude.[19,40] In particular, Rubisco displays extremely limited variation in $k_{cat,C}$ ($\sigma^*$ = 1.5), especially as compared to other enzymes for which multiple $k_{cat}$ measurements are available (median $\sigma^*$ = 6.9, Figure S5).[41]

Form I and Form II enzymes differ most in their $S_{C/O}$, $k_{cat,C}$ and $K_C$ values, which can be seen by comparing the median of blue and yellow distributions in Figure S4A. This observation is consistent with the notion that carboxylation parameters are under the most stringent selection.[22] In addition to analyzing variation between Rubisco isoforms, the extended dataset can be used to see variation in kinetic parameters between Rubiscos from different hosts. Figure S6 shows that different host physiologies are associated with characteristic carboxylation kinetics, with $C_3$ plant Rubiscos being more $CO_2$-specific than cyanobacterial Rubiscos, for example.

#### Correlation and Regression Analyses

Certain model Rubiscos are characterized frequently. For example we found 12 independent measurements of the model Rubisco from spinach and 10 for the model cyanobacterial Rubisco from *Synechococcus* PCC 6301. We used the median measured value in correlation and regression analyses to avoid bias towards frequently-characterized Rubisco variants. Once multiple measurements were merged, we used performed correlation and regression analyses on a log scale to investigate coupling between Rubisco parameters. The reasoning for log-scale analyses is discussed in detail below, but is based on our expectation (rooted in transition state theory) that relationships between kinetic parameters will have exponential form. An exponential relationship of the form $Y = a X^k$ is called a power law. A power law relation produces a linear relationship on a log-log plot: $\log(Y) = \log(a) + m \log(X)$. The slope of the linear fit corresponds to the power law exponent and the Y-intercept corresponds to the exponential prefactor.

Regressions were performed using total least squares (TLS), which is sometimes also called orthogonal distance regression (ODR). TLS regression is applicable to cases where there is error associated with both X and Y variables, which is the case here because both variables are experimental measurements derived from fits to data. However, $R^2$ values of TLS fits do not convey the explained fraction of Y axis variance and are therefore not easily interpreted. As such, we report the degree of correlation as Pearson correlation, R. $10^4$-fold bootstrapping was used to estimate 95% confidence intervals for R, power-law exponents and prefactors. In each iteration of the bootstrap, data were subsampled to 90% with replacement. The 95% confidence interval on R gives an indication of the robustness of the underlying power-law correlation. Results of correlation and regression analyses are discussed in the main text.

#### Use of Log-Scale Correlations

Throughout the main text we presented log-scale correlations and regressions to investigate tradeoffs between Rubisco kinetic parameters. For reasons described here we strongly prefer to investigate these questions on a logarithmic scale. Nevertheless, linear correlations and scatter plots are in Figure S7. Figure S7A shows that pairwise linear-scale correlations are qualitatively similar to the log scale correlations in Figure 4. As in log-scale, the strongest linear scale correlation is between the catalytic efficiencies for carboxylation and oxygenation, $k_{cat,C}/K_C$ and $k_{cat,O}/K_O$ (R = 0.90). Plotting the catalytic efficiencies $k_{cat,C}/K_C$ against $k_{cat,O}/K_O$ (Figure S7D) shows that robust log-scale correlation is recapitulated on a linear scale, as would be expected when $S_{C/O}$ is roughly constant.

Log-scale correlations are appropriate for two reasons. First, the proposed tradeoff mechanisms invoke coupling of TS barrier heights. Additive tradeoffs between TS barriers should manifest as power law correlations between rate constants, as discussed below. Second, linear regression on a logarithmic scale is more robust to multiplicative error, which is common in biochemical measurements and likely present in this case.

If two TS barriers, $\Delta G_1$ and $\Delta G_2$, trade off with each other, the tradeoff will manifest as some coupling between the barrier heights. We assume that the coupling is linear:

$$\Delta G_1 + m\Delta G_2 = b$$
$$exp\ (\Delta G_1) = exp\ (-\Delta G_2)^m \times exp\ (b)$$

Where *m* and *b* are positive constants and ΔG values are in units of RT (R being the universal gas constant and T the temperature in Kelvin) so that constants can be omitted for simplicity. Transition state theory posits that kinetic constants $k_1$ and $k_2$ are proportional to exponential functions of the barrier height:

$$k_1 \propto exp\ (-\Delta G_1)$$
$$k_2 \propto exp\ (-\Delta G_2)$$

It follows from these equations

$$\frac{1}{c_1 k_1} = exp\ (b) \times (c_2 k_2)^m$$
$$k_1 = c' {k_2}^{-m}$$

Here $c_1$ and $c_2$ are constants of proportionality, and $c' = (e^{-b}\ c_1^{-m}) / c_2$ is a constant as well. Therefore, we expect to find a negative power law correlation with exponent -*m* between two kinetic constants when there is a negative linear relationship between their respective TS barrier heights (with slope -*m*). Given particular assumptions elaborated below, measurable Rubisco kinetic parameters can also be treated as exponentially related to effective TS barriers. It is therefore preferable to perform correlation analysis on a log scale so as to interpret correlations between kinetic parameters are related to the microscopic kinetics of Rubisco.

Log-scale regressions are also appropriate when measurement errors are multiplicative. That is, when the error scales with the measured quantity as implied by saying “within 10% error” or similar. Multiplicative error is common in many experimental settings and error in Rubisco kinetic measurements appears to scale with measured values as well (Figure S3). Multiplicative error suggests that errors are normally distributed on a log scale. Least-squares regression assumes that error is normally distributed, so it is sensible to perform regression on a log scale when errors are multiplicative.

#### Principal Component Analysis

We repeated the PCA analysis of Savir et al. 2010 to test if the principal axis of variation is similar in our larger dataset. As in Savir et al. 2010, PCA was performed on a 4 dimensional space of parameters that uniquely determine net carbon fixation: [$K_C$, $k_{cat,C}$, $S_{C/O}$, $K_C$ / $K_O$].[22] Analysis was restricted to those Rubiscos for which $S_{C/O}$, $K_C$, $k_{cat,C}$, and $K_O$ measurements were available from the same reference. This dataset contains roughly 250 more Rubiscos than that of Savir et al. 2010.[22] The data were log-transformed and Z-score normalized before PCA. Normalization and PCA was performed using the Python sklearn package.

Focusing on Form I Rubiscos shows that the principal axis remains mostly unchanged by the addition of new data. PC1 = [ 0.47 0.52 -0.48 0.53] over the Form I data in the extended dataset as compared to PC1 = [0.52 0.50 -0.48 0.51] over the 15 FI data points reported in Savir et al 2010.[22] These vectors are quite similar, with a cosine distance of 0.004 implying a roughly 5° angle between them. However, The proportions associated with the principal components (% of variance explained) are now [70% 12% 11% 5%] as compared to [91%, 6%, 2%, 0.6%] in the smaller dataset of Savir et al. 2010, indicating that there is substantially more variation in the extended dataset and that a one-dimensional approximation may not be appropriate (Figure S8).

#### Measurement Error and Systematic Error in Rubisco Measurements

In the main text and above we discuss correlations in the extended dataset and highlight reduction in pairwise correlations in comparison to previous analyses of smaller datasets.[22,25] Rubisco assays are challenging to perform and variation in measurements across labs is expected. Some of the spread in the data may come from systematic differences between labs and assay methods. Rubisco activation state (i.e. degree of carbamylation) may differ between methods and preparations, for example.[28] As such, we were careful to review each study's methods and record experimental error when reported and filtered a small number of suspect measurements, as discussed above.

Weakened correlations could be due to measurement errors in a larger dataset. We find this explanation unlikely for several reasons. First, several correlations remain statistically significant (Figures 4-7), including the very strong power-law correlation between $k_{cat,C}/K_C$ and $k_{cat,O}/K_O$ (Figure 7). Despite lab-to-lab variation, we multiple measurements of the same Rubisco are broadly consistent (Figure S4B) and Rubiscos from similar organisms (e.g. $C_3$ or $C_4$ plants) have very similar $S_{C/O}$ values that are clearly distinct from other groups (Figure 5A and 7B). Finally, experimental error is reported for > 85% of measured values and the average reported error is < 15% of the mean value for all directly measured parameters (Figure S3).

The fact that Rubisco requires activation by carbamylation of an active site lysine complicates Rubisco assays and might be a source of systematic experimental error due to overestimation of the number of active sites.[28,42] Purified Rubisco is typically incubated with excess $CO_2$ before enzymatic assays to ensure full activation.[38] Still, the activation state may vary across preparations and assay conditions. As such, it is now standard to quantify the number of active sites by binding to the stoichiometric inhibitor, CABP.[30,38] This method is considered reliable and enables reproducible $k_{cat,C}$ measurements. Recent work also demonstrates that standard Rubisco assays produce results that are quantitatively consistent with a mass-spectrometric method directly measuring carboxylation and oxygenation reactions.[43] The data making up the extended dataset are generally recent and mostly use CABP to count active sites, which substantially ameliorates concerns about systematic underestimation of $k_{cat,C}$ and $k_{cat,O}$.

Residual analysis is another means of checking for systematic bias in a dataset (Figure S9). Overall, residual distributions are roughly symmetric about 0 and are quite similar to those from the smaller dataset of Savir et al.,[22] suggesting that measurement errors are similarly distributed in the extended dataset. The $k_{cat,C}$-$K_C$ fit offers an interesting counter-example. Outlying measurements are clearly biased to lie beneath the best-fit power-law (Figure S9B). This result implies one of two possibilities: either some outlying measurements underestimate $k_{cat,C}$ and/or $K_C$, or some natural Rubiscos are strictly worse than others on these axes. Outliers are mostly from non-green algae and bacteria and include several recent high-quality measurements of diatom Rubiscos.[44] These organisms have $CO_2$ concentrating mechanisms, which is expected to relax selection for $CO_2$ affinity and could explain why these measurements lie beneath the fit line.

#### Correlations with $k_{cat,O}$

An intriguing suggestion of Savir et al. 2010 is that $k_{cat,O}$ might vary independently of other kinetic parameters.[22] It is reasonable to assume that carboxylation is selected for. If $k_{cat,O}$ is not under strong selection, lack of correlation between $k_{cat,O}$ and other measured kinetic parameters might imply that selection is required for correlation to emerge ("selection within limits" model in Figure 2B).[22] However, the model described in the main text (Figures 7 and 8) invokes coupling between the kinetics of oxygenation and carboxylation to explain the limited variation in $S_{C/O}$ and strong power-law correlation between $k_{cat,C}/K_C$ and $k_{cat,O}/K_O$. As such, there are conflicting views as to whether $k_{cat,O}$ should correlate with other Rubisco kinetic parameters.

Figure S10 evaluates log-scale $k_{cat,O}$ correlations in the extended dataset. Focusing on parameters that are directly measured, $k_{cat,O}$ appears to correlate modestly with $k_{cat,C}$ and $K_C$. Power-law correlation between $k_{cat,C}/K_C$ and $k_{cat,O}/K_O$ implies that there should be some correlation between oxygenation and carboxylation parameters, so this result is not unexpected. Indeed, $k_{cat,O}$ correlates well with $k_{cat,C}/K_C$ on a log scale ($R > 0.5$). Future research will hopefully resolve correlations between carboxylation and oxygenation parameters by direct measurement of $k_{cat,O}$ and $K_O$ via mass spectrometry and other methods.[43]

### Relationship Between Tradeoff Models and the Microscopic Kinetics of Rubisco

#### Derivation of Rubisco Kinetic Equations

Figure S1 diagrams the microscopic kinetic scheme for Rubisco carboxylation and oxygenation following the nomenclature of Cummins et al. 2018.[45] A complete derivation of the Michaelis-Menten type rate law for Rubisco carboxylation and oxygenation is given in the supplement of that reference. This derivation makes only one assumption: that product release (of carboxylation and oxygenation products both) is irreversible, i.e. that $k_{10}$, $k_{15}$ = 0 (Figure S1).

#### Catalytic Efficiencies ($k_{cat}/K_M$) are Related to the First Effective Barrier to Carboxylation and Oxygenation

One central result from the derivation[45] is that the catalytic efficiencies for carboxylation and oxygenation can be expressed as:

$$\frac{k_{cat,C}}{K_C} = \frac{k_{cat,C} K_R k_5}{k_{cat,C} + \gamma_C k_6}$$

$$\frac{k_{cat,O}}{K_O} = \frac{k_{cat,O} K_R k_{11}}{k_{cat,O} + \gamma_O k_{12}}$$

Where $\gamma_C$and $\gamma_O$ are defined as

$$\gamma_C = \frac{k_3 k_8 + k_3 k_9}{k_3 k_7 + k_3 k_8 + k_3 k_9 + k_7 k_9}$$

$$\gamma_O = \frac{k_3 k_{14} + k_3 k_{15}}{k_3 k_{13} + k_3 k_{14} + k_3 k_{15} + k_{13} k_{15}}$$

$k_5$ is the rate constant for $CO_2$ association with the Rubisco-enediolate complex (ER*) and $k_6$ represents $CO_2$ dissociation from the ERC complex. Similarly, $k_{11}$ is the rate of $O_2$ association ER* complex and $k_{12}$ represents $O_2$ dissociation from the ERO complex.$K_R = \frac{k_3}{k_3 + k_4}$is related to the equilibrium fraction of the Rubisco-RuBP complex in the enediolate form ($K_E = \frac{ER^*}{ER} = \frac{k_3}{k_4}$; $K_R = \frac{K_E}{1+K_E}$). In a commentary on Cummins et al. 2018,[45] Tcherkez et al. 2018[46] summarize experimental evidence that decarboxylation and deoxygenation rates are small in comparison to carboxylation and oxygenation, respectively. As a result, it is usually assumed decarboxylation and deoxygenation are small in comparison to the forward reactions (i.e. $k_{cat,C} \gg \gamma_C k_6$and $k_{cat,O} \gg \gamma_O k_{12}$)[22,23,25,46] to derive that

$$\frac{k_{cat,C}}{K_C} \approx K_R k_5$$

$$\frac{k_{cat,O}}{K_O} \approx K_R k_{11}$$

There remains some disagreement about the validity of this assumption.[45,46] Nonetheless, under these assumptions $k_{cat,C}/K_C$ and $k_{cat,O}/K_O$ are determined by enolization and gas addition rates alone.

The relationship between catalytic efficiencies and gas addition rates can be understood intuitively as follows. $k_{cat}/K_M$ is the slope of the Michaelis-Menten relationship ($v = \frac{k_{cat}[S]}{[S]+K_M}$) in the limit of low substrate ($[S] \ll K_M$ implies $v \approx \frac{k_{cat}}{K_M}[S]$). In the case of Rubisco, $k_{cat,C}/K_C$ represents the balance of $CO_2$ association and dissociation in the subsaturation limit. If $CO_2$ dissociation is negligible then $k_{cat,C}/K_C$ reflects association alone and should therefore be proportional to the microscopic association rate.

Savir et al. 2010 apply transition state theory to obtain *effective* barriers to enolization and gas addition:

$$\frac{k_{cat,C}}{K_C} \propto exp\,(-\Delta G_{1,C})$$

$$\frac{k_{cat,O}}{K_O} \propto exp\,(-\Delta G_{1,O})$$

Here the effective barriers to enolization and gas addition $-\Delta G_{1,C}$ and $-\Delta G_{1,O}$ in units of RT.[22] Note that the left hand side of these expressions have units of $s^{-1}$ $\mu M^{-1}$ meaning that the true barrier heights will depend on the $CO_2$ and $O_2$ concentrations.[25] The $CO_2$ concentration near Rubisco can vary between organisms, especially because some have CCMs and others do not. $C_3$ plants lack a $CO_2$-concentrating mechanism and so their Rubiscos experience roughly the same $CO_2$ concentration. Measurements of the $CO_2$ partial pressure in $C_3$ plant leaves typically lie between 65-85% of ambient,[47,48] meaning that the soluble $CO_2$ concentration likely varies by at most 30% within this group. For this reason we highlighted the $C_3$ plants in describing Figure 6B in the main text.

#### $k_{cat,C}$ is Related to the Second Effective Barrier for Carboxylation

Following the notation of Cummins et al. 2018,[45]

$$k_{cat,C} = \frac{k_3 k_7 k_9}{k_3 k_7 + k_3 k_8 + k_3 k_9 + k_7 k_9}$$

$k_3$ is the only term in this expression that relates to enolization. If hydration and bond cleavage are rate limiting, as is often assumed,[22,49] then the $k_7 k_9$ term should be small compared to the other terms in the denominator as $k_7$ and $k_9$ are the rate constants associated with those two steps (Figure S1). In that limit, $k_3$ cancels and

$$k_{cat,C} \approx \frac{k_7 k_9}{k_7 + k_8 + k_9}$$

$k_{cat,C}$ is therefore roughly independent of enolization ($k_3$) if hydration and cleavage are rate-limiting for carboxylation. As such, $k_{cat,C}$ can be related to an *effective* barrier to hydration and cleavage of the 6-carbon carboxyketone intermediate.

$$k_{cat,C} \propto exp\,(-\Delta G_{2,C})$$

Here $-\Delta G_{2,C}$ is expressed in units of RT, as above. A similar result can be obtained for $k_{cat,O}$ but the oxygenation mechanism is poorly understood[23] and such an expression is not required to interpret our main-text figures.

#### Interpretation of Power-Law Correlation Between $k_{cat,C}/K_C$ and $k_{cat,O}/K_O$

In the main text we documented very restricted and stereotyped variation in $S_{C/O}$ and argued that this forces strong power-law correlation between $k_{cat,C}/K_C$ and $k_{cat,O}/K_O$ (Figure 7). This can be understood by examining the definition of $S_{C/O}$:

$$S_{C/O} \equiv (\frac{k_{cat,C}}{K_C})/(\frac{k_{cat,O}}{K_O})$$

$$(\frac{k_{cat,C}}{K_C}) = S_{C/O} \times (\frac{k_{cat,O}}{K_O})$$

$$log\ (\frac{k_{cat,C}}{K_C}) = log\ (S_{C/O}) + log\ (\frac{k_{cat,O}}{K_O})$$

So if $S_{C/O}$ is roughly constant, then it is the Y-intercept of a power law relationship between $k_{cat,C}/K_C$ and $k_{cat,O}/K_O$ with exponent 1.0.

If decarboxylation and deoxygenation rates are roughly 0 ($k_6, k_{12} \approx 0$), the catalytic efficiencies can be expressed as functions of enolization and gas addition alone, as described above. Substituting into the expression above gives:

$$log\ (K_R) + log\ (k_5) = log\ (S_{C/O}) + log\ (K_R) + log\ (k_{11})$$

The enolization equilibrium constant $K_R$ cancels and $S_{C/O}$ is roughly constant (Figure 7B), so positively correlated variation in $k_5$ and $k_{11}$ must drive the power law correlation between $k_{cat,C}/K_C$ and $k_{cat,O}/K_O$ shown in Figure 7C if all these assumptions are valid.

#### Derivation of the Active Site Gating Model

The model diagrammed in Figure 8 envisions a Rubisco-RuBP complex that is found in either a "reactive" or "unreactive" state. Note that these states might be mapped onto the catalytic mechanism drawn in Figure S1 as the RuBP must be RuBP is isomerized to an enediolate for either reaction to proceed.[27,28] As such, the "reactive" state is likely related to the ER* complex. Our model is coarse grained, however, and the mapping to the Rubisco mechanism diagrammed is likely not direct. For example, conformational rearrangements (e.g. closure of loop 6) are required for carboxylation.[28,50] These rearrangements should be accounted for in any definition of "reactive" and "unreactive" states of the Rubisco active site.

In the "reactive" state both $CO_2$ and $O_2$ can and react with the enediolate of RuBP with their intrinsic reactivities, denoted by barrier heights $\Delta G^*_{1,C}$and $\Delta G^*_{1,O}$. The fractional occupancy of the reactive state is denoted $\phi$. As discussed above, this factor $\phi$ could arise solely from enolization of RuBP as well as other factors, so long as these factors affect $CO_2$ and $O_2$ equally. Whatever factors determine the reactivity of the active site, we assume that they are fast-equilibrating.

Given these assumptions,

$$k_{cat,C}/K_C \propto \phi\ exp\ (-\Delta G^*_{1,C}/RT)$$

$$k_{cat,O}/K_O \propto \phi \; exp\;(-\Delta G^*{}_{1,O}/RT)$$

Where the constants of proportionality are assumed to be the same in both cases. If enolization is the only factor determining reactivity, then $\phi = K_R = \frac{K_E}{1+K_E}$and $\phi^{-1} = e^{\Delta G_E/RT} + 1$. As above, ΔG values are in units of RT for simplicity. $\phi$ cancels when we calculate $S_{C/O}$.

$$S_{C/O} \; = \; (k_{cat,C}/K_C)/(k_{cat,O}/K_O) \quad = exp\;(\Delta G^*{}_{1,O} - \Delta G^*{}_{1,C})$$

If the intrinsic reactivities $\Delta G^*{}_{1,C}$and $\Delta G^*{}_{1,O}$ are organism independent, then $S_{C/O}$ should be constant. Taking the log of both sides we can also derive the expected power-law correlations.

$$log\;(S_{C/O}) = log\;(k_{cat,C}/K_C\;\;) - log\;(k_{cat,O}/K_O) = \Delta G^*{}_{1,O} - \Delta G^*{}_{1,C}$$

At first glance it would seem that $\phi = 1$would be the best possible outcome for all Rubiscos because this would yield the same constant $S_{C/O}$ but maximize $k_{cat,C}/K_C$. However, this intuition stems from a misunderstanding of $S_{C/O}$. $S_{C/O}$ is not trivially related to the net rate of carboxylation. Rather, $S_{C/O}$ is the slope of $R_C/R_O$ when it is plotted against $[CO_2]/[O_2]$ in the limit of low $CO_2$ and $O_2$ concentrations. Simple manipulation of the equations for $R_C$ and $R_O$ gives

$$R_C = k_{cat,C}[CO_2]([CO_2] + \; K_C \; + \; K_C[O_2]/K_O)^{-1}$$

$$R_O = k_{cat,O}[O_2]([O_2] + \; K_O \; + \; K_O[CO_2]/K_C)^{-1}$$

$$R_C/R_O = \frac{k_{cat,C}[CO_2]}{k_{cat,O}[O_2]}\frac{([O_2] \; + \; K_O + K_O[CO_2]/K_C)}{([CO_2] + K_C + K_C[O_2]/K_O)}$$

In the limit where $[CO_2] \ll \;\; K_C$and $[O_2] \ll \;\; K_O$we can simplify as follows: $[CO_2] + K_O \approx \;\; K_O$and $K_O[CO_2]/K_C \ll \;\; K_O$. In this limit

$$R_C/R_O \approx \frac{(k_{cat,C}/K_C)}{(k_{cat,O}/K_O)} \times \frac{[CO_2]}{[O_2]} = S_{C/O} \times \frac{[CO_2]}{[O_2]}$$

However, this only applies in the limit of low $CO_2$ and $O_2$ concentrations. The full relationship between $S_{C/O}$ and $R_C/R_O$ is not linear and, moreover, $S_{C/O}$ does not uniquely set the net carboxylation rate *f* = $R_C$ - 0.5 $R_O$. value of  As such, readers should not assume that higher $S_{C/O}$ being better for carboxylation.

#### Implications of the Active Site Gating Model

The implication of the data presented in Figures 7-8 and the model described above is that $\phi$ varies within the various Rubisco groups, perhaps by varying the barrier to RuBP enolization ($\Delta G_E$), which does not appear in the expression $S_{C/O} = \frac{k_5}{k_{11}}$ derived above. Varying $\phi$ will cause a proportional increase in both $k_{cat,C}/K_C$ and $k_{cat,O}/K_O$ which can produce the power law correlation shown in Figure 7C. The model does not explain why there is spread in $k_{cat,C}/K_C$ and $k_{cat,O}/K_O$, e.g. why certain $C_3$ plants have higher $k_{cat,C}/K_C$ values than others. We suspect that these variations are related to organismal growth conditions like temperature, humidity and salinity,[51] but we do not currently have sufficient data to investigate these hypotheses.

According to power law derived from our coarse-grained model, $S_{C/O}$ should be approximately constant. However, $S_{C/O}$ varies about tenfold between Form I and Form II Rubiscos and 3-4 fold among the Form I enzymes. When we examine Rubiscos isolated from hosts belonging to the same physiological group - e.g. $C_3$ or $C_4$ plants - they do display a characteristic and roughly constant $S_{C/O}$ (Figure 7B).

Restricted variation in $S_{C/O}$ implies $\Delta G^*_{1,O} - \Delta G^*_{1,C}$ varies little within the groupings depicted in Figure 7B-C. The fact that $C_3$ plants, $C_4$ plants and cyanobacterial Rubiscos assume characteristic $S_{C/O}$ values could be explained as evolutionary optimization of $\Delta G^*_{1,O} - \Delta G^*_{1,C}$ given prevailing $CO_2$ and $O_2$ concentrations. However, since the dataset is no longer approximately one-dimensional, it is not straightforward to make this argument quantitative.[22] We hope future work will investigate the degree to which Rubisco is optimized to suit environmental conditions and explain why particular Rubiscos assume higher $k_{cat,C}/K_C$ values than others.

#### Characterized Rubisco Mutants Do Not Exceed Wild-Type Tradeoffs

Various schemes have been used to select Rubisco mutants for characterization. We extracted data for ≈40 mutant Rubiscos. We were able to calculate both catalytic efficiencies $k_{cat,C}/K_C$ and $k_{cat,O}/K_O$ for 15 mutants. These mutants are predominantly 1-3 amino acid substitutions to wild-type enzymes, but there is also kinetic data on chimeric rubiscos[52] and reconstructed ancestral sequences.[26] Cyanobacterial FI[24,26,52–57] and proteobacterial FII[58,59] Rubiscos are the most commonly mutagenized. Figure S11 plots $k_{cat,C}/K_C$ against $k_{cat,O}/K_O$ for these mutant enzymes along with matching wild-type (WT) enzymes for comparison. Most mutants are "worse" than WT in that $k_{cat,C}/K_C$ is lower than would be expected based on the WT trend. Several of the cyanobacterial Rubisco mutants are strictly less efficient than the WT enzymes, with $k_{cat,C}/K_C$ and $k_{cat,O}/K_O$ more than tenfold lower than WT values (Figure S11).

### Supplementary Tables

***Table S1:*** *Description of supplementary datasets provided with this manuscript.*

| Dataset Name | Description |
|---|---|
| Dataset S1 | Full source data including all annotations. |
| Dataset S2 | Filtered and merged source data including inference of measurement error for primary and derived parameters like catalytic efficiencies. |
| Dataset S3 | Data from the BRENDA database used to generate Figure S4. |
| Dataset S4 | An unfiltered version of Dataset S2 that includes mutant data used to generate Figure S11. |

***Table S2:*** *Description of the columns in Dataset S1, S2 and S4. Column names ending in "_SD" are reported standard deviations for their respective values. Column names ending in "_95CI_low" and "_95CI_high" are the inferred low and high ends of the 95% confidence interval for this value, respectively. * $k_{cat,O}$ is very rarely measured directly. It is inferred from other parameters in nearly all cases. Therefore values in Dataset S1 should also be considered inferences even though they were extracted from primary literature in most cases.*

| Column Label | Description |
|---|---|
| species | Name of the host species. Attempted consistency across multiple measurements of the same Rubisco. |
| identifier | A unique identifier of this row, usually of the form species_reference |
| primary | 1 if the reference is the primary reference for this measurement. Otherwise 0. |
| mutant | 1 if this is a mutant Rubisco, e.g. point mutants, hybrids or inferred ancestral sequences. Otherwise 0. |
| heterologous | 0 if this Rubisco was purified from the native host. Otherwise the name of the organism used for heterologous expression. |
| KC | Measured Michaelis constant for CO2 in μM units. |
| vC | Measured $k_{cat,C}$ in $s^{-1}$ units. |
| S | Measured $S_{C/O}$. |
| KO | Measured Michaelis constant for O2 in μM units. |
| vO | *Inferred $k_{cat,O}$ in $s^{-1}$ units. |
| vO_reported | *Original reported $k_{cat,O}$ - nearly always an inference too. |
| kon_C | Inferred catalytic efficiency for $CO_2$ $k_{cat,C}/K_C$ in $s^{-1}$ $\mu M^{-1}$ units |
| kon_O | Inferred catalytic efficiency for $O_2$ $k_{cat,O}/K_O$ in $s^{-1}$ $\mu M^{-1}$ units |
| KRuBP | Michaelis constant for RuBP in μM |
| temp_C | Celsius temperature at which the measurement was taken. |
| pH | pH at which the measurement was taken. |
| isoform | Rubisco isoform - Form I, II, II/III and III are encoded as 1, 2, 2_3 and 3. |
| variant | For Form I only. Denotes if they are Form IA, B, C or D. Inferred from species. |
| taxonomy | A taxonomic tag used to group Rubiscos, e.g. in Figures 6-8. |
| note | Any notes about this value. |
| short_ref | A short name for this reference. |
| pmid_or_doi | Identifier of this reference. Pubmed ID when available, DOI otherwise. |
| citation | Full citation for this reference. |

### Supplementary Figures

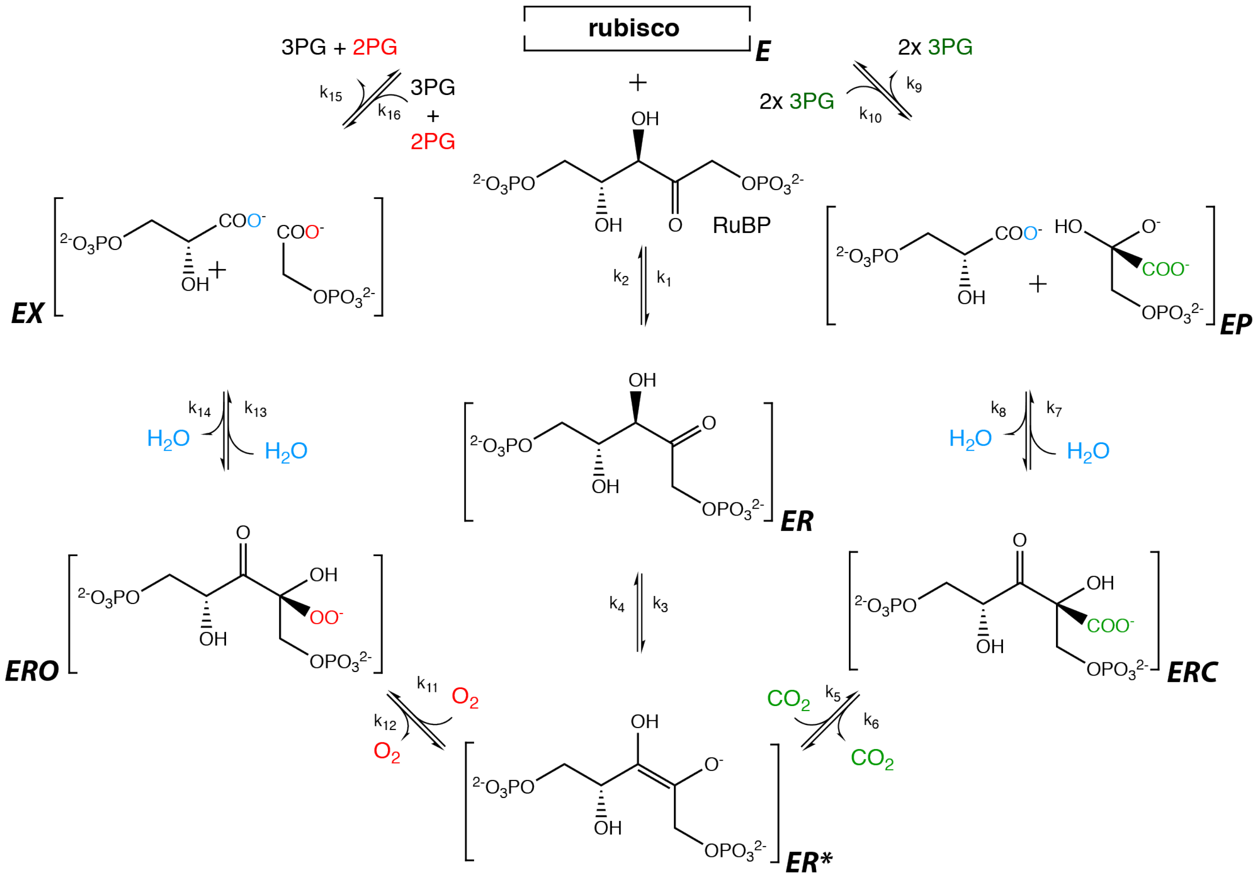


***Figure S1:*** *The mechanism of RuBisCO following the nomenclature of Cummins et al. 2018 [45]. RuBisCO must be carbamylated and then bind Mg2+ before it becomes catalytically active, after which it processes three substrates - ribulose bisphosphate (RuBP), $CO_2$ and $O_2$. Rubisco can bind the five-carbon RuBP before or after activation. RuBP binding before activation strongly inhibits activity and RuBP is removed by a catalytic chaperone called Rubisco activase in many organisms [5,6]. The complete reactions of carboxylation and oxygenation take place through a stepwise mechanism [25,27,28]. RuBP binds first forming a complex (ER) with the activated form of the enzyme (E), followed by enolization of RuBP (ER*) which allows binding and further processing of $CO_2$ or $O_2$. When $CO_2$ binds the ER* complex the ERC complex is formed while $O_2$ binding leads to formation of the ERO enzyme-substrate complex. Hydration and cleavage of the ERC complex leads to the formation of two enzyme-bound 3-phosphoglycerate molecules (3PG) in the EP state, each of which have 3 carbon atoms. Oxygenation proceeds through analogous steps except that the products contain 5 carbon atoms in total instead of 6 because no carbon was added. Hydration and cleavage of the ERO complex produces one enzyme-bound 3PG and one 2-phosphoglycolate (2PG) in the EX state. 2PG has two carbon atoms and is not part of the CBB cycle. As such it must be recycled through a photorespiratory pathway to avoid the accumulation of 2PG and also the loss of two carbons [11]. Atoms originating from free $CO_2$ and $O_2$ are shown in green and red respectively. The oxygen atom originating from water is shown in blue.*

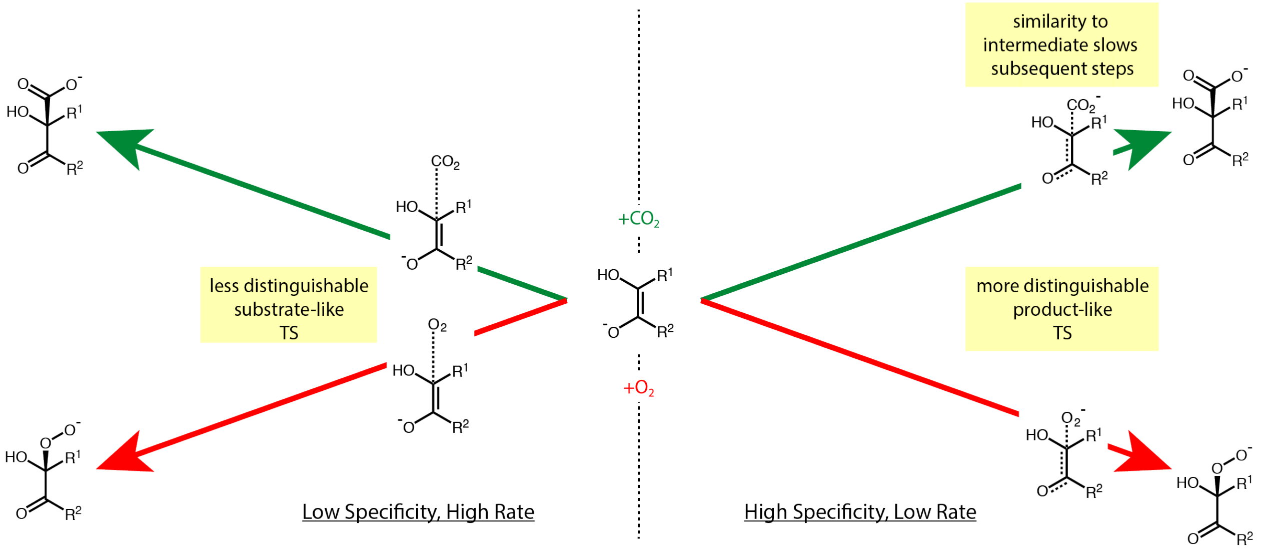


***Figure S2:*** *Visual explanation of the tradeoff as proposed by* [25]*. In this model, it is assumed that Rubisco discriminates between $CO_2$ and $O_2$ at the level of the first transition state. The need for selectivity in the presence of high $O_2$ or low $CO_2$ concentrations promotes a "late" carboxylation transition state (green trajectory in the right half of the figure) which more resembles the carboxylate of the carboxyketone intermediate. The developing carboxylate of a late TS is maximally discriminable from the peroxyacid of the first oxygenation TS (red trajectory). However, in this model, stabilization of a late carboxylation TS forces tight binding of the carboxyketone carboxylation intermediate (following Hammond's postulate) and slows the downstream rate-limiting hydration and cleavage of the intermediate. Though this model is motivated by the need for selectivity, note that the tradeoff implied by the model is between initial binding of $CO_2$ ($k_{cat,C}/K_C$) and subsequent processing of the carboxylation intermediate ($k_{cat,C}$). That is, the tradeoff is strictly independent of the $O_2$ concentration and affects only those kinetic parameters related to carboxylation.*

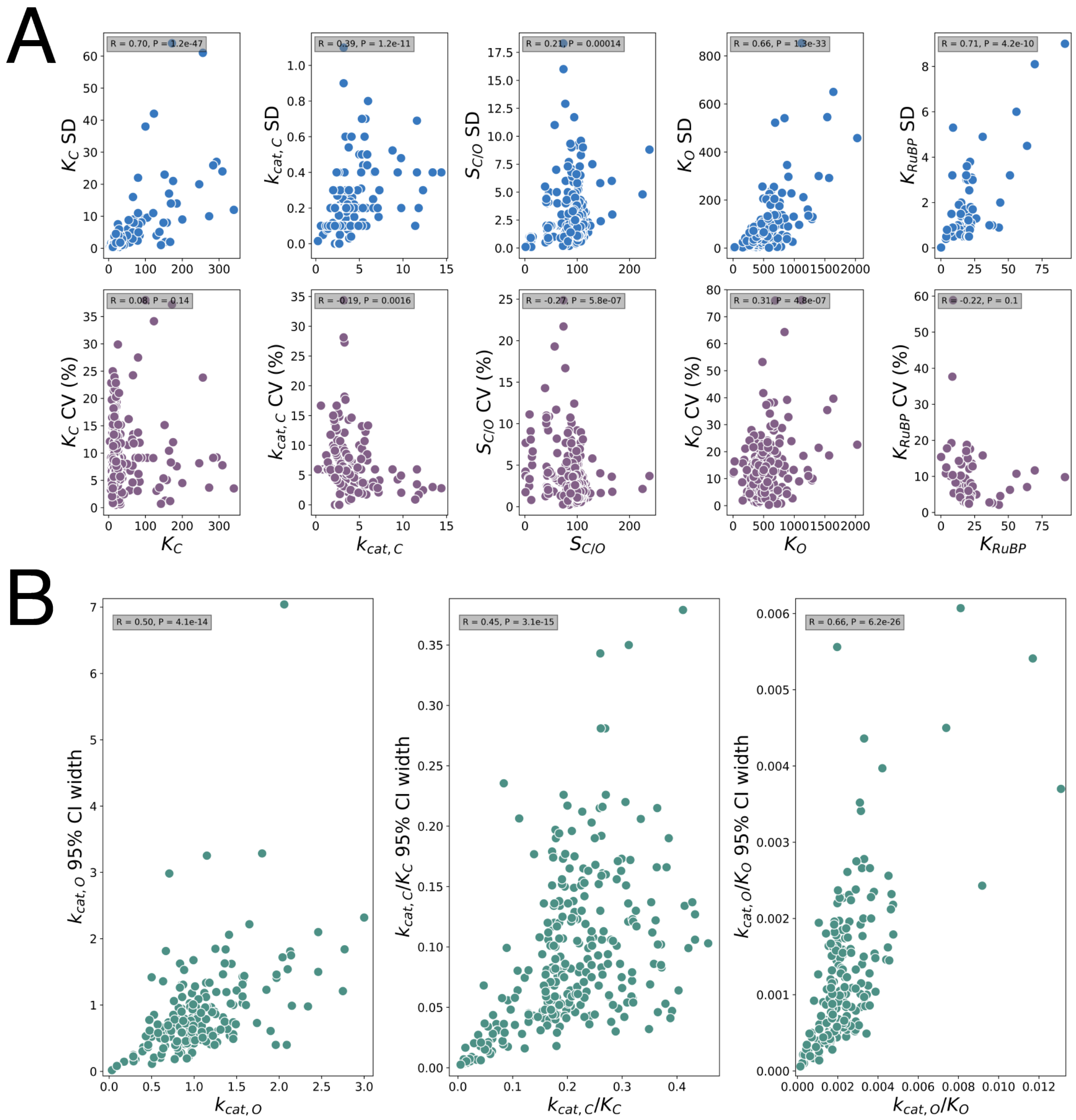


***Figure S3:*** *Measurement error scales with measured values. This justifies the use of regression on a logarithmic scale. (A) Reported measurement error scales with the mean value for all directly measured parameters, consistent with multiplicative error (top row). The second row shows that the coefficient of variation (CV, standard deviation divided by the mean value) does not depend as strongly on the measured value, as expected for multiplicative error. (B) kcat,O, kcat,C/KC and kcat,O/KO are inferred from the other measurements and 95% confidence intervals (CIs) are calculated by bootstrapping as described in the Methods section. CIs for these values also appear to scale with the measured value, supporting log-scale regression for derived values as well.*

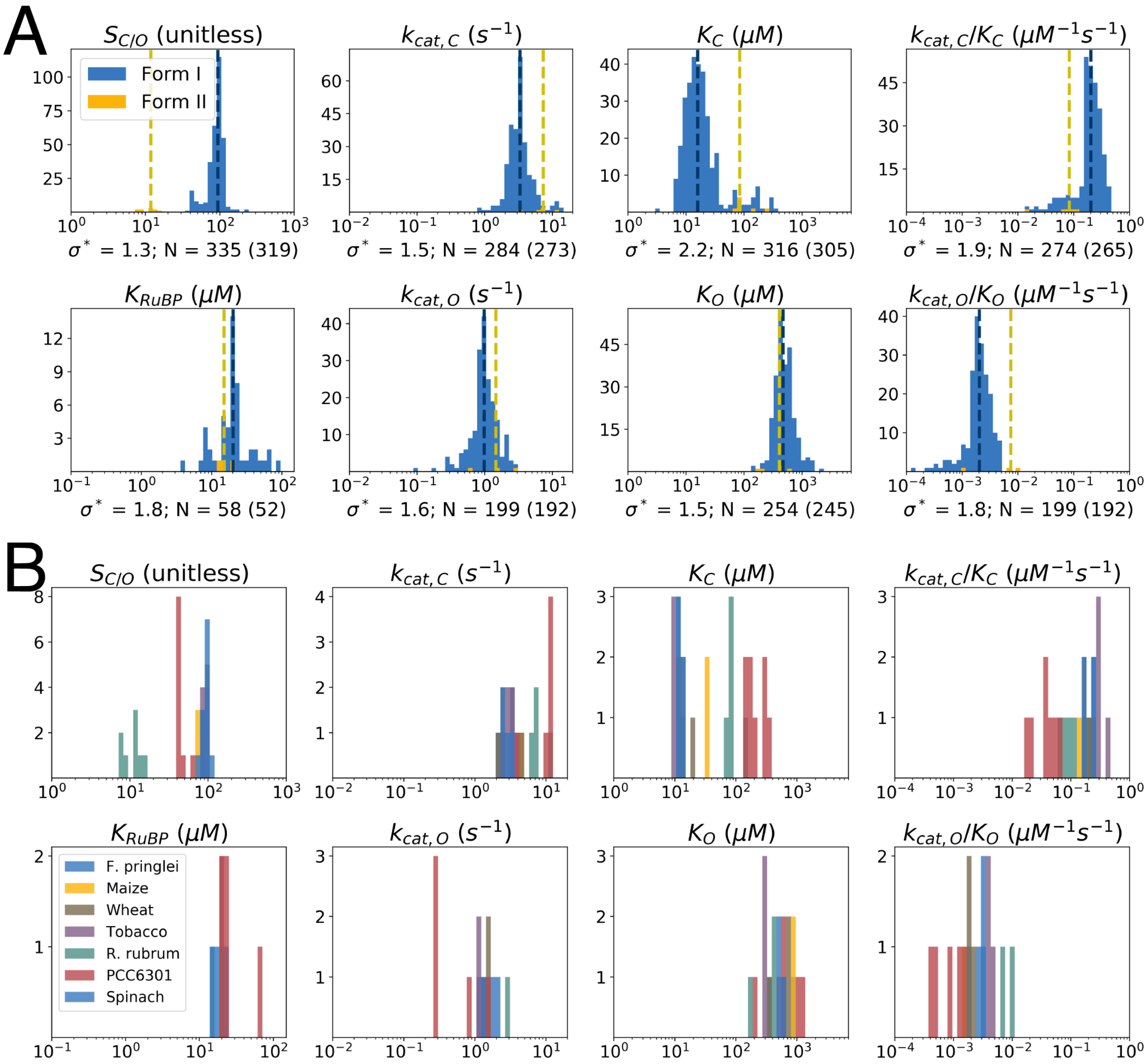


***Figure S4:*** *Histograms of Rubisco kinetic parameters. Kinetic parameter values are on the X-axis (log scale) and number of data points is on the Y-axis. (A) Parameter distributions for all Form I and Form II Rubiscos in the extended dataset. Form I distributions are in blue and Form II distributions are in yellow. Other Rubisco isoforms are omitted because data is scant. Dashed lines give the median value. N is the number of Form I values and $\sigma$* is the multiplicative standard deviation of Form I values. (B) Distributions for the seven Rubiscos with > 3 measurements each. In the main text figures we used median values to represent species with multiple measurements. Multiple measurements of the same Rubisco are broadly consistent, though early $k_{cat,C}$ measurements of the Synechococcus PCC6301 Rubisco are roughly fivefold lower than recent measurements, likely due to improved quantitation of the number of active sites. We found N=4 distinct measurements of Form I Rubisco from F. pringlei, N=4 of maize (Z. mays), N=7 of wheat (T. aestivum), N=8 of tobacco (N. tabacum), N=10 of S. elongatus PCC 6301, N=12 of Spinach (S. oleracea), and N=9 of the Form II Rubisco from R. rubrum.*

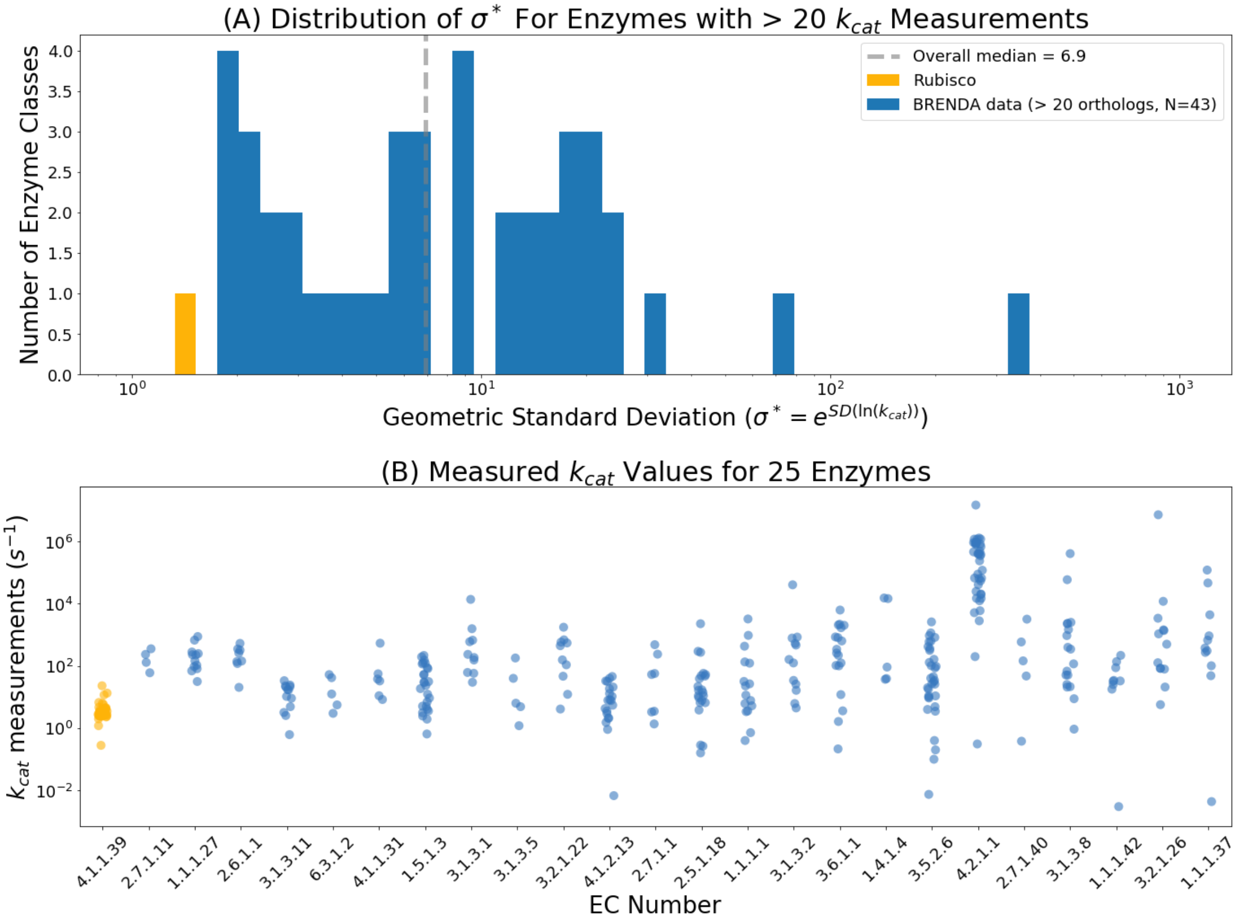


***Figure S5:*** *Rubisco displays extremely limited $k_{cat,C}$ variability as compared to other enzymes. $k_{cat}$ data for wild-type enzymes other than Rubisco was drawn from the BRENDA database.[60] In cases where multiple measurements of the same enzyme from the same organism were found, the median value was used. This procedure produced 240 $k_{cat,C}$ values for Rubisco with a multiplicative standard deviation $\sigma^* = 1.5$. When the underlying distribution is log-normal, a value of $\sigma^* = 2.0$ connotes that the central 68% of values are within two-fold of the mean. (A) Rubisco is an outlier among enzymes for which >20 kcat measurements are available, displaying fivefold less $k_{cat}$ variability than the median enzyme (median $\sigma^* = 6.9$). (B) $k_{cat}$ data for the 25 enzyme classes with the most data, ordered from least-to-most multiplicative variability (left to right). Rubisco is in yellow. Roughly fivefold more data is available for Rubisco than any other enzyme, but it displays the least variation in $k_{cat}$ of any enzyme in the dataset.*

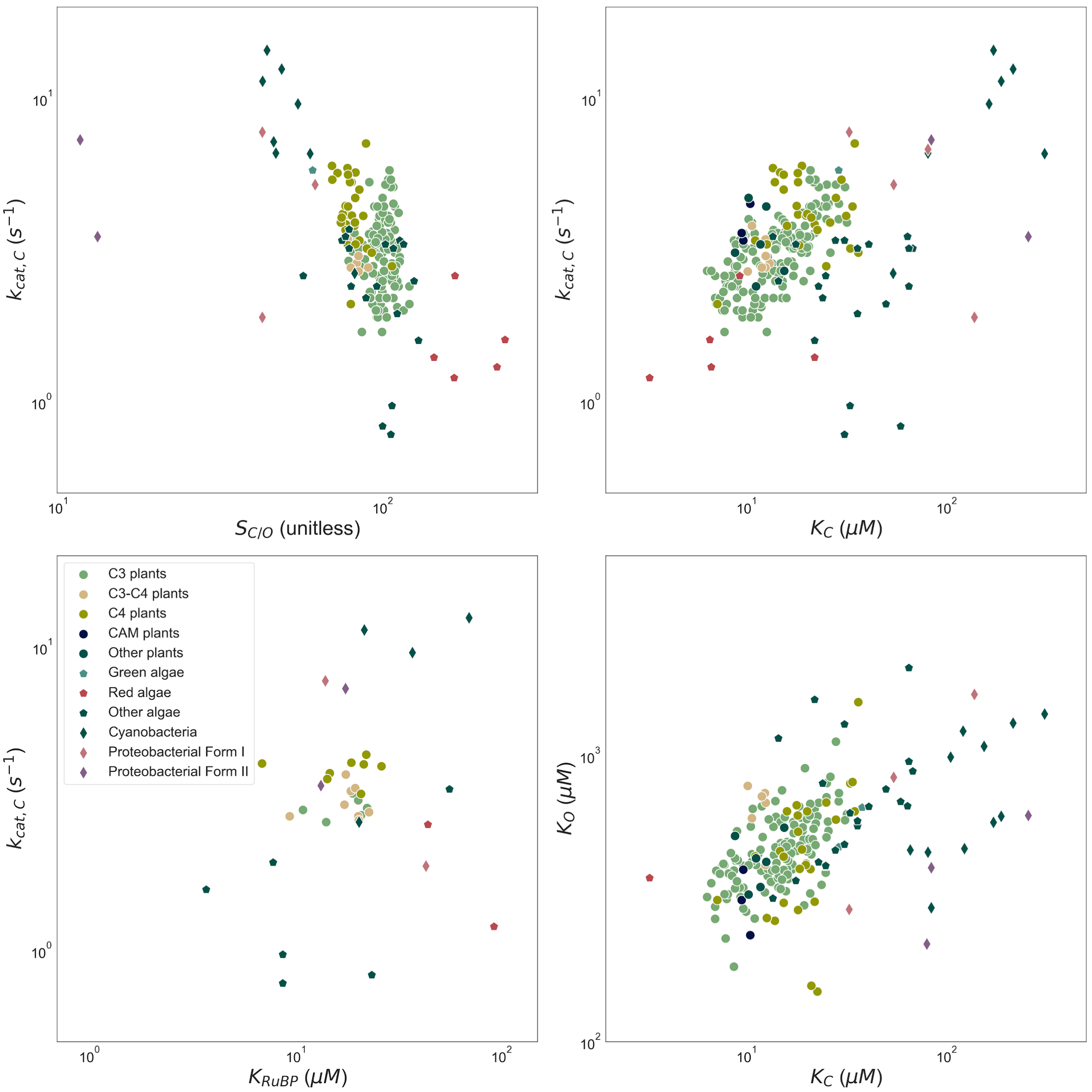


***Figure S6:*** *Rubisco kinetic parameters by host type. Main text figures merge multiple classes of hosts into a single category ("other") for clarity and simplicity. This figure reproduces main-text plots while retaining more granular distinctions between organismal physiology and phylogeny. Importantly, it is challenging to define a uniform taxonomy for autotrophs because physiological characteristics (e.g. CCMs) are not monophyletic. The ad-hoc taxonomy used here and in supplementary datasets is imperfect but useful for representing characteristic differences in Rubisco kinetics. Cyanobacterial rubiscos generally have the highest reported* $k_{cat,C}$ *values and red algal Rubiscos have characteristically high* $CO_2$ *specificity. Different host physiologies generally occupy distinct regions of the* $k_{cat,C}$ *vs.* $S_{C/O}$ *plot (top-left).*[25] *However, as documented in the main text, these broad trends do not produce a very strong correlation, which is what we would expect if Rubisco was "perfectly optimized" with respect to a tradeoff between* $k_{cat,C}$ *and* $S_{C/O}$.

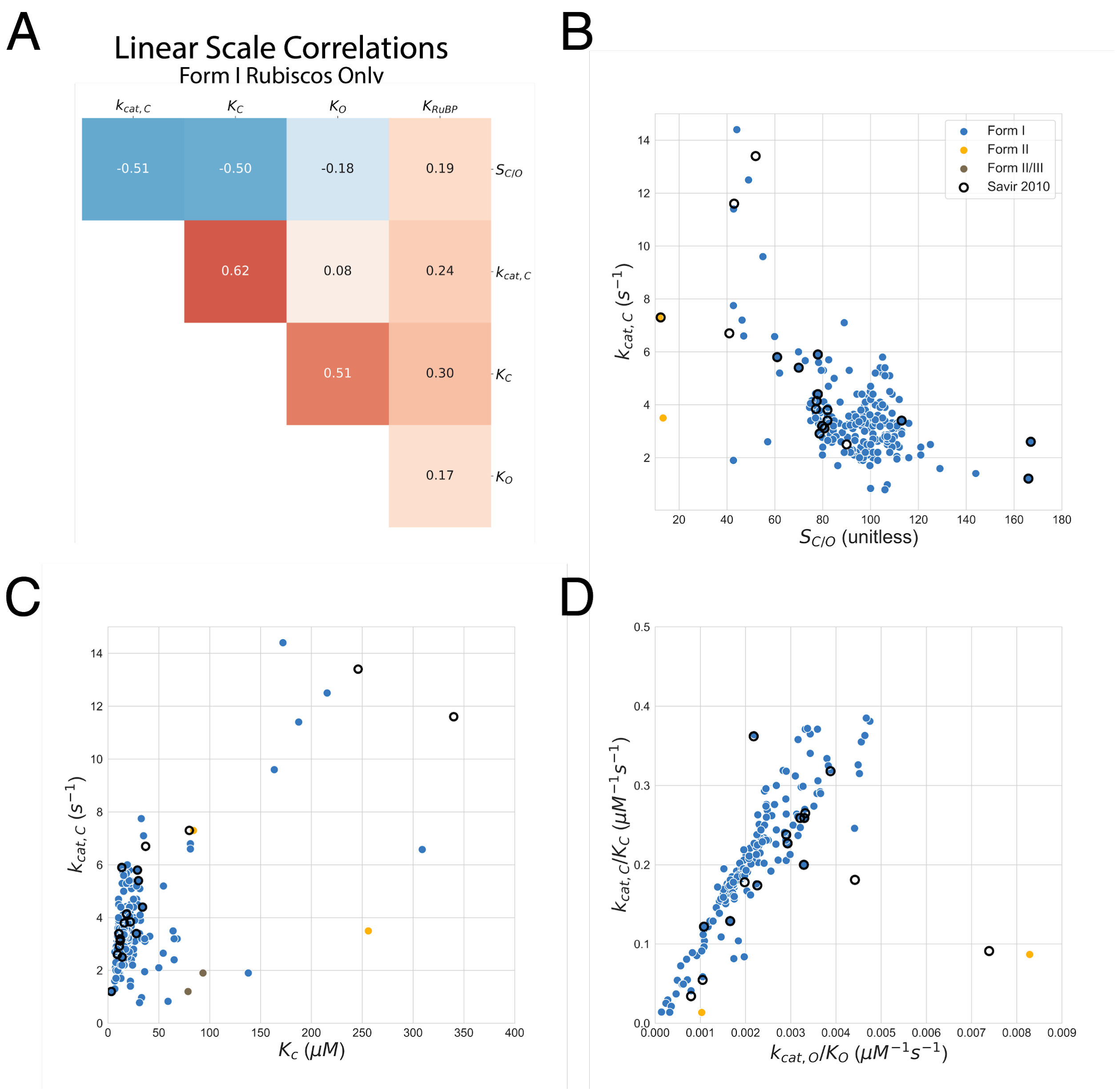


***Figure S7:*** *Central correlation figures in linear scale. (A) A heatmap of pairwise linear-scale correlations. (B) Linear-scale scatter plot of $k_{cat,C}$ against $S_{C/O}$ shows that the data has very limited dynamic range and correlations are determined by a small number of outliers. (C) Linear scatter plot of $k_{cat,C}$ against $K_C$. (D) Linear scatter plot of catalytic efficiencies $k_{cat,C}/K_C$ against $k_{cat,O}/K_O$ shows that robust log-scale correlation is recapitulated on a linear scale, as would be expected for a power-law exponent of 1.0.*

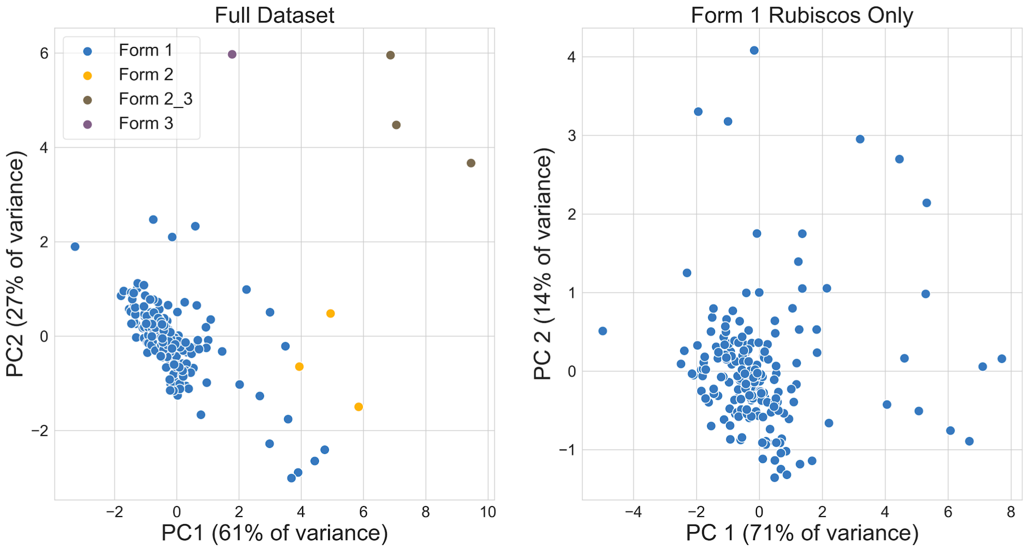


***Figure S8:*** *Principal components analysis of the extended dataset. Data were normalized before PCA in order to ensure that the very different absolute values of Rubisco kinetic parameters did not bias analysis. Left panel gives the projection onto the first two principal components (PCs) determined by the subset of the extended dataset for which sufficient data was available, i.e. for which the vector [$k_{cat,C}$, $K_C$, $S_{C/O}$, $K_C$ / $K_O$] could be calculated. The first two PCs separate the Rubisco isoforms from each other. When PCA is restricted only to Form I RuBisCOs, as in the right panel, there is still substantially more variation in Rubisco kinetic parameters than described in previous work, where PC1 accounted for roughly 90% of the variation in Rubisco kinetics. In this setting, Rubisco was described as evolving within a roughly one-dimensional landscape.[22] PC1 now explains less of the variance (71%) of the variance, so Rubisco kinetics cannot be described accurately as varying in a one-dimensional landscape.*

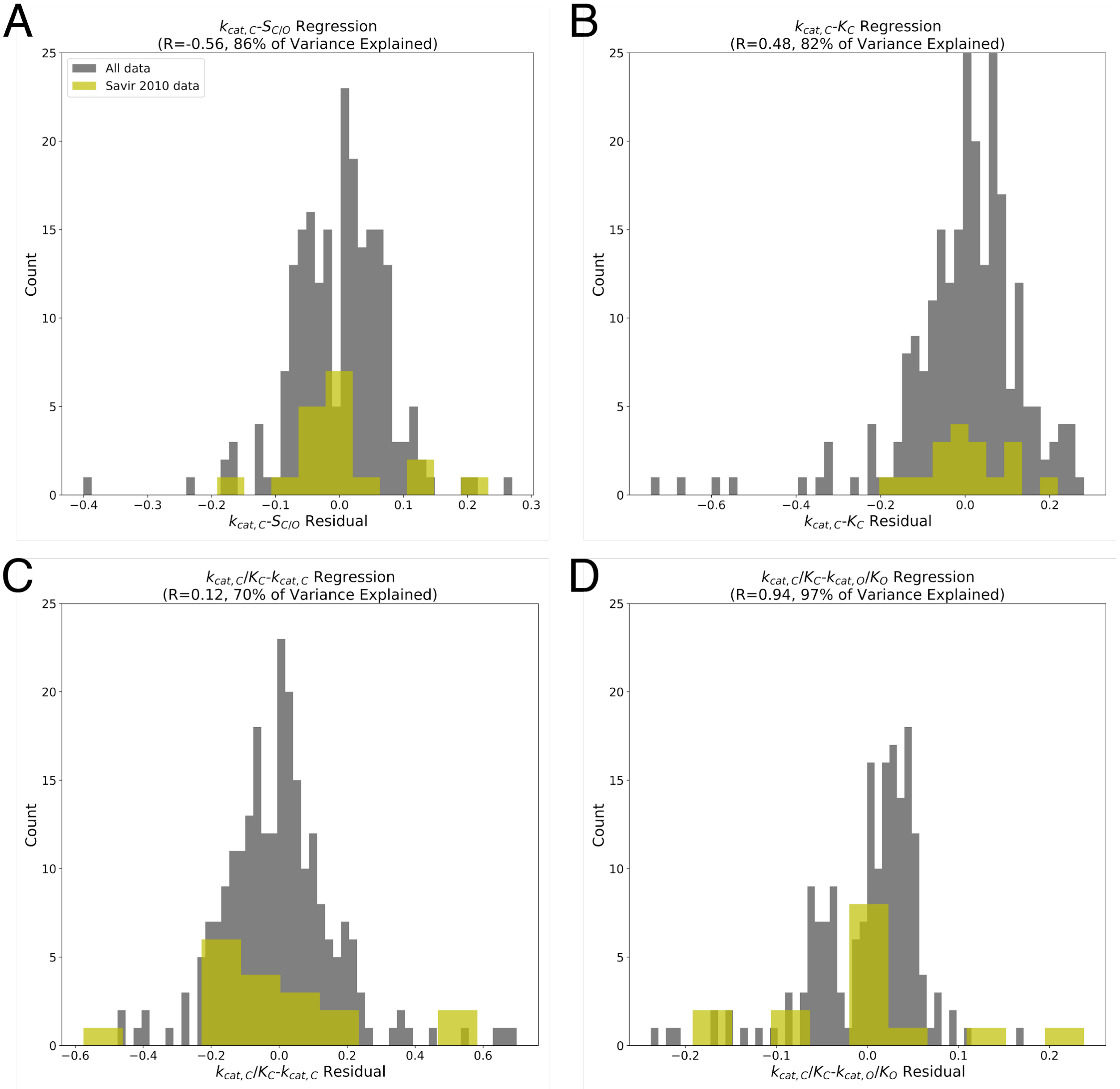


***Figure S9:*** *Residuals for total least squares fits of FI Rubiscos. Two dimensional data are rotated so that the axis of greatest variation (first principal component) is on the X-axis. Residuals are calculated as distance to the X-axis for the whole dataset (grey) and the subset from Savir et al. 2010 (yellow). Panel (A) gives residuals for $k_{cat,C}$-$S_{C/O}$ fit; (B) Residuals for $k_{cat,C}$-$K_C$ fit; (C) Residuals of ($k_{cat,C}/K_C$)-$k_{cat,c}$ fit; (D) Residuals of ($k_{cat,C}/K_C$)-($k_{cat,O}/K_O$) fit. (B) Outliers in the full dataset tend to lie beneath the $k_{cat,C}$-$K_C$ regression line, suggesting that a tradeoff between these variables follows the "selection within limits" model (Figure 2B). Outliers include recent measurements of Rubiscos from diatoms.[44] In panel (D), in contrast, addition of tenfold more data has not altered the distribution of residuals appreciably. This supports a "mechanistic coupling" between ($k_{cat,C}/K_C$) and ($k_{cat,O}/K_O$) as diagrammed in Figure 2A.*

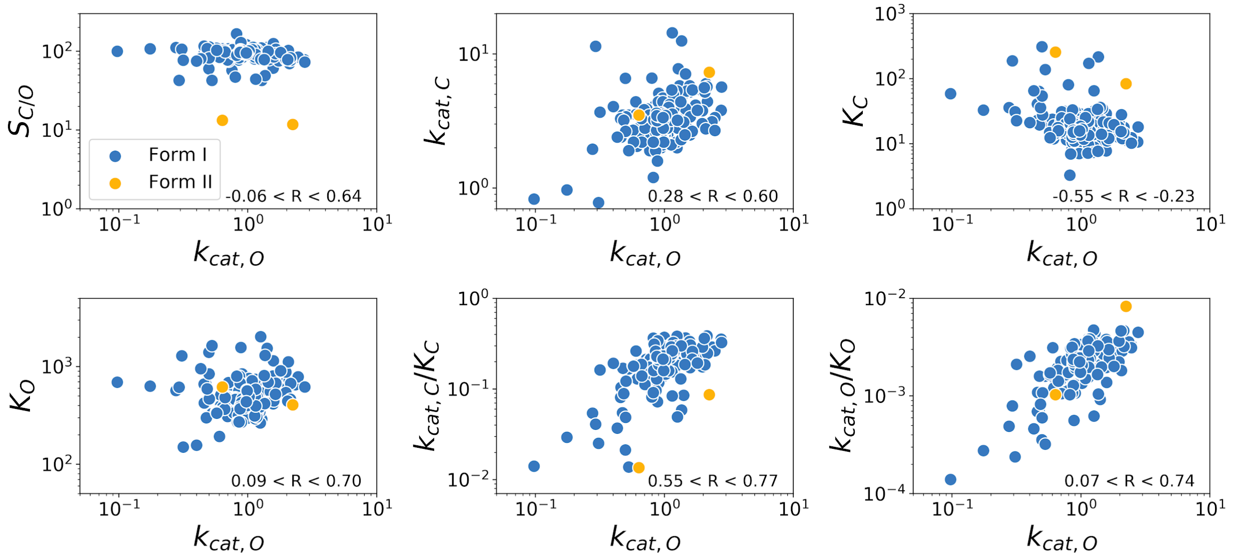


***Figure S10:*** *Correlations between $k_{cat,O}$ and other kinetic parameters. $S_{C/O}$ is unitless, $k_{cat}$ values have $s^{-1}$ units and $K_M$ values have μM units. Form I Rubiscos are in blue and FIIs are in yellow. A 95% confidence interval on the Pearson correlation R among Form I enzymes by 1000-fold bootstrapping and indicates that the $k_{cat,O}$ may be correlated with $k_{cat,C}$, $K_C$, and $k_{cat,C}/K_C$.*

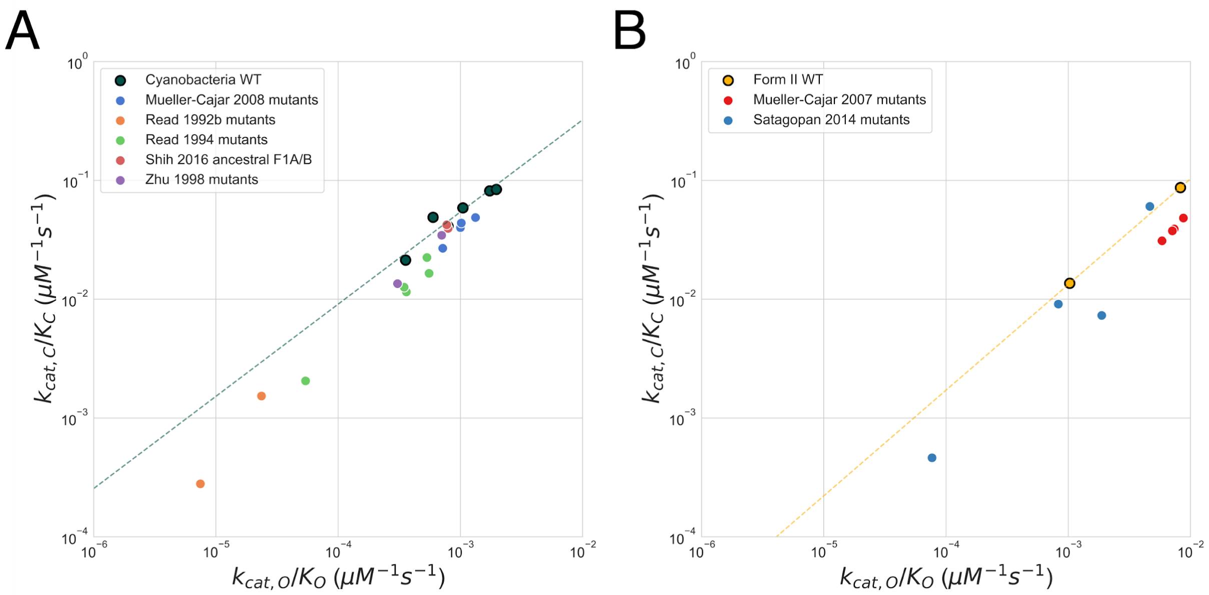


***Figure S11:*** *Mutant cyanobacterial Form I and proteobacterial Form II Rubiscos do not exceed tradeoffs calculated from counterpart wild-type enzymes. (A) Scatter plot of $k_{cat,C}/K_C$ against $k_{cat,O}/K_O$ for mutant cyanobacterial Form I enzymes. The dashed green line gives the best-fit regression line determined by the WT cyanobacterial enzymes. Note that several of the mutants from Read & Tabita references[52,54] have tenfold lower catalytic efficiency towards both $CO_2$ and $O_2$ than all WT cyanobacterial enzymes. (B) Mutant Form II Rubiscos from proteobacteria. The dashed yellow line gives the best-fit regression line determined by 3 WT enzymes. In both panels the best fit is determined by < 10 data points and should not be considered high-confidence. Rather the data here are given to suggest that mutant Rubiscos investigated so far do not surpass the catalytic efficiency tradeoff determined for comparable WT enzymes.*
